# Transcription-coupled changes in higher-order genomic structure and transcription hub viscosity prolong enhancer-promoter connectivity

**DOI:** 10.1101/2023.11.27.568629

**Authors:** Hiroaki Ohishi, Soya Shinkai, Hitoshi Owada, Takeru Fujii, Kazufumi Hosoda, Shuichi Onami, Takashi Yamamoto, Yasuyuki Ohkawa, Hiroshi Ochiai

**Affiliations:** Division of Gene Expression Dynamics, Medical Institute of Bioregulation, Kyushu University; Fukuoka, Japan; Laboratory for Developmental Dynamics, RIKEN Center for Biosystems Dynamics Research; Kobe, Japan; Graduate School of Integrated Sciences for Life, Hiroshima University; Higashi-Hiroshima, Japan; Division of Transcriptomics, Medical Institute of Bioregulation, Kyushu University; Fukuoka, Japan; Ansanga Lab; Suita, Japan

## Abstract

The orchestration of our genes heavily relies on a coordinated communication between enhancers and promoters, yet how this dynamic interplay remains elusive while transcription is active. Here, we investigated enhancer-promoter (E-P) interactions and relative to transcriptional bursting in mouse embryonic stem cells using sequential DNA/RNA/immunofluorescence (IF)-FISH analyses and computational simulations. Our data reveal that the active state of specific genes is characterized by higher-order genomic structures and local condensates of transcriptional regulatory factors, leading to an elevation in local viscosity that highly stabilizes the duration of E-P interactions. Our study underscores the pivotal role of viscosity in transcriptional dynamics and paves the way for a more nuanced understanding of gene-specific regulatory mechanisms.

## Introduction

The transcription of cell-type-specific genes is governed by enhancer regions that often are distant from the gene promoters. Enhancer regions are rich in binding sites for transcription factors that are expressed in a cell type specific manner. For an enhancer to coordinate the cell-specific information instructed via transcription factors to a cognate promoter, it must physically be located near a promoter region. For example, a clear correlation between enhancer-promoter (E-P) proximity and transcriptional activity has been demonstrated by live-cell imaging in Drosophila^1^. Conversely, several cases have been reported in which no apparent correlation between E-P proximity and transcriptional activity has been observed^2–6^. These findings are compatible with the transcription hub model, which posits that E-P communication occurs through transcriptional regulatory factor condensates and does not necessitate direct contact between the E-P pair^7^. Meanwhile, recent Micro-C analyses have uncovered clear interactions between E-P pairs just downstream of the transcription start site, at the location of the +1 nucleosome^8^. However, the temporal aspects of these interactions remain undetermined.

Imaging and sequencing-based analyses have revealed that transcription is a dynamic process characterized by stochastic switches between active states, where RNA is continuously synthesized, and inactive states, where little to no synthesis occurs. This phenomenon, universally observed across various species, cell types, and genes, is commonly referred to as transcriptional bursting^9,10^. Transcriptional bursting plays a critical role in regulating both the level and cell-to-cell heterogeneity in gene expression. Recent findings indicate that during the active state, coactivators such as BRD4—which binds to acetylated histones at active enhancer regions—and the large subunit of RNA polymerase II (RPB1), form condensates in the spatial proximity of the gene locus^11^. Additionally, the gene region exhibits slower mobility in the active state^11,12^. These observations suggest that higher-order genomic structures, including E-P interactions, may undergo dynamic changes that regulate transcriptional dynamics. However, significant knowledge gaps remain in our understanding of the relationship between these dynamically changing E-P interactions and transcriptional dynamics. To bridge these gaps, our study employed both sequential (seq) DNA/RNA/immunofluorescence (IF) fluorescence *in situ* hybridization (seq-DNA/RNA/IF-FISH) analyses and computational polymer simulations. The seq-DNA/RNA/IF-FISH analyses allowed us to visualize transcriptional activity states, higher-order genomic structures, transcriptional regulatory factors, and post-translational modifications (PTMs) at the level of single alleles within individual mouse embryonic stem (ES) cells.

## Results

### Multiplexed imaging of chromatin structure and transcriptional activity

To explore the relationship between transcriptional dynamics and higher-order genomic structure, we utilized imaging-based seq-DNA/RNA/IF-FISH analyses. This innovative technology combines seq-RNA-FISH, seq-DNA-FISH, and seq-immunofluorescence FISH (seq-IF-FISH) on the same sample, thereby offering a comprehensive view of transcriptional activity, higher-order genomic structures, and transcriptional regulatory factors/PTMs at single-allele resolution within individual cells (Fig. 1a, Supplementary Fig. 1)^13^.

**Fig. 1.**
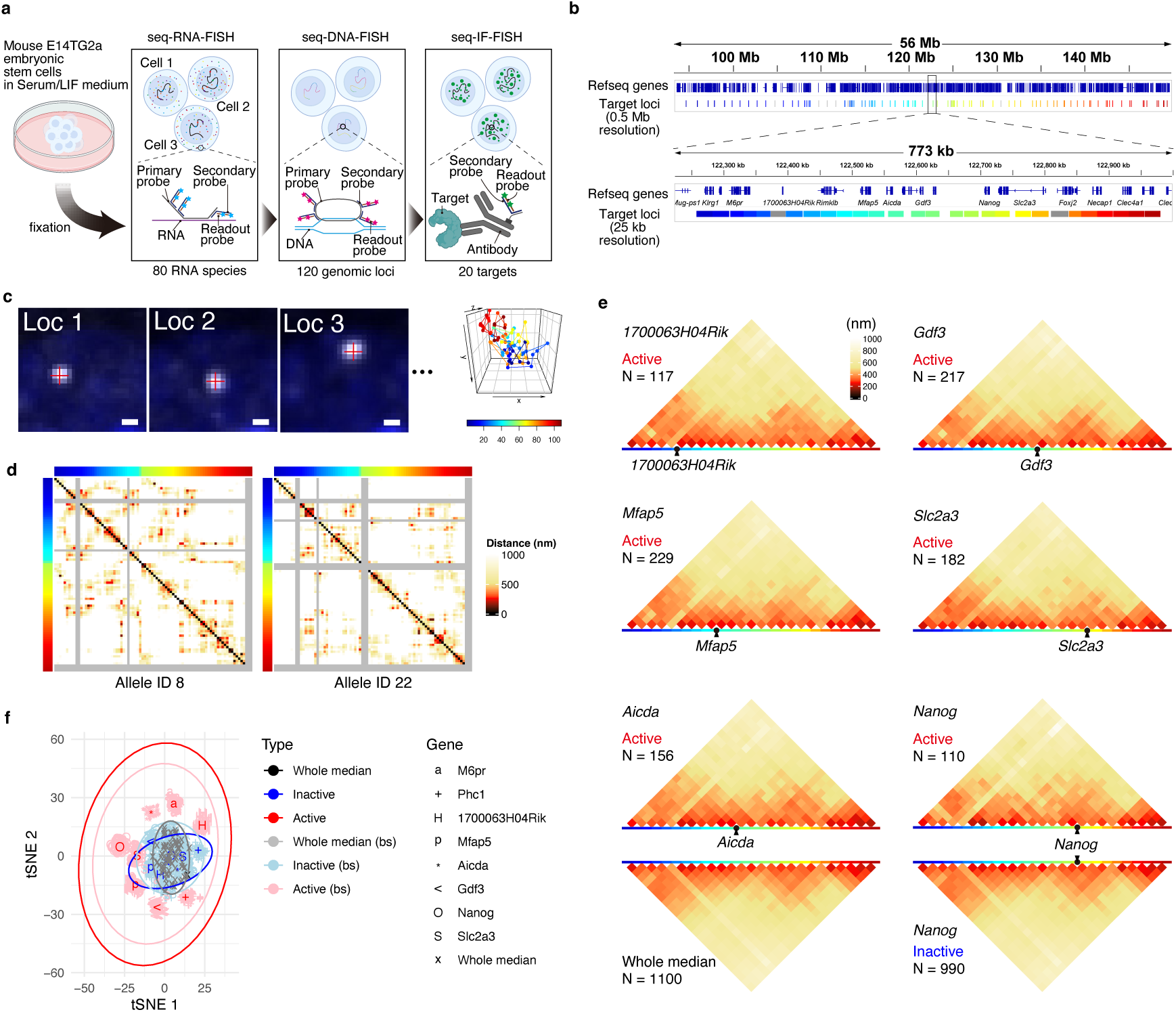
Transcriptionally active state-specific higher-order genomic structures. **a** Schematic representation of seq-DNA/RNA/IF-FISH analysis. **b** Target genomic regions in seq-DNA-FISH analysis. The upper panel represents target regions at a 0.5 Mb resolution, while the lower panel shows them at a 25 kb resolution. The gray regions designate target regions excluded from detailed analysis due to quality control checks. **c** Representative examples of fluorescent spots detected in seq-DNA-FISH imaging. The scale bar in the image represents 500 nm. The color code at the bottom of the right panel corresponds to the target genomic regions in **b** at 0.5 Mb resolution. **d** Distance matrices for specific alleles, revealing noticeable differences between alleles. The gray regions indicate areas where fluorescent spots were not detected. Color codes on the top and left margins correspond to the 0.5 Mb resolution target genomic regions in **b**. **e** Median distance matrices classified by the active/inactive states of specific genes. The number of samples is indicated within the Figure. Color codes correspond to the 25 kb resolution target genomic regions in **b**. “Whole median” refers to the median value calculated from all the data. **f** t-SNE analysis of median distance matrices, classified by whether the gene is in an active or inactive state. The term “Whole median” refers to the median distance matrix calculated without dividing the data into active or inactive states for a specific gene. Using bootstrap methods (bs), we randomly sampled from the alleles, allowing for duplicates, and created median distance matrices the same number of times as the sample size *N*. The results from 100 bootstrap samples are shown.

We focused our investigation on the *Nanog* gene, a key regulator of pluripotency, in mouse ES cells cultured in a medium containing leukemia inhibitory factor (LIF) and serum (serum/LIF medium). In this condition, where pluripotency is maintained, transcriptional bursting at the *Nanog* locus has been previously observed^14^. 4C-seq analyses of mouse ES cells have shown that the *Nanog* locus can interact with genomic regions spanning several to tens of megabases^15^ (Supplementary Fig. 2a). Although these distal interactions are relatively weak compared to the proximal regions of *Nanog* and occur across multiple A compartments (Supplementary Fig. 2a), the functional consequences of these interactions on transcriptional bursting remain unclear. To elucidate the functional consequences of these distal interactions, we subjected genomic regions—both including and excluding these domains—to seq-DNA-FISH analyses at intervals of approximately 0.5 Mb. In addition, we subjected regions for seq-DNA-FISH analysis at 25 kb intervals within an approximately 750 kb region surrounding the *Nanog* locus (Fig. 1b, Supplementary Fig. 2a). This 750 kb region largely comprises a singular topologically associating domain (TAD) (Supplementary Fig. 2). In total, we analyzed 120 genomic loci spanning approximately 60 Mb for seq-DNA-FISH analysis (Figs. 1a,b).

Within the 60 Mb range targeted for seq-DNA-FISH analyses, we also chose 80 highly expressed genes in mouse ES cells as targets for seq-RNA-FISH (Supplementary Fig. 2a, see Methods). Furthermore, to correlate the epigenetic state, transcription, and higher-order genomic structure, we targeted 20 types of entities that include transcriptional regulatory factors and histone PTMs. These are notably enriched in *Nanog*-interacting regions and were investigated using seq-IF-FISH (see subsequent sections) (Supplementary Movie 1).

From seq-DNA-FISH analyses, we obtained coordinates of foci for each gene region, enabling distance measurements between genomic loci (Fig. 1c). Some targeted regions had low detection frequencies, leading us to exclude them from our subsequent analyses (Supplementary Fig. 3a, see Methods). Consequently, we proceeded with 109 remaining loci for in-depth analysis (Supplementary Fig. 3a). We performed two individual seq-FISH experiments and acquired data from 611 cells and 1,100 alleles in Replicate 1, and 335 cells and 580 alleles in Replicate 2. We generated two-dimensional distance matrices from the 3D coordinates, revealing extensive heterogeneity in higher-order genomic structures, consistent with previous reports^16,17^ (Fig. 1d). A high inverse correlation between the median distance matrices and Hi-C contact frequency data was observed (Supplementary Fig. 3b). Additionally, a high degree of correlation was found between replicates when comparing median distance matrices (Supplementary Fig. 3c, d), indicating the reliability of the acquired data. Unless otherwise specified, the results presented herein are based on data from Replicate 1.

### Transcriptionally active state-specific higher-order genomic structures

Utilizing our seq-RNA-FISH data, we visualized individual RNA molecules. In regions where transcription is actively occurring, numerous nascent RNAs accumulate, resulting in the detection of bright fluorescence spots. In this study, bright spots detected near specific alleles, which were more intense than those from single RNA molecules, were identified as ’transcriptional spots’. We used these spots as an indication that the gene is in an active state (see Methods). Upon calculating the proportion of transcriptionally active alleles for each gene, we found that only about 10% of the genes (8 out of 80) exhibited an average active allele proportion exceeding 20% (Supplementary Fig. 3e). This observation is in accord with prior studies that suggest most genes exhibit transcriptional bursting^10,18^. We subsequently categorized genome structures based on their transcriptional activity states and examined their median distance matrices. Using data at 0.5 Mb resolution, we discerned that the differences between transcriptional activity states were relatively minor. This observation suggests that the infrequent interaction between A compartments does not exert a strong influence on transcriptional dynamics (Supplementary Fig. 3f). When focusing on data at the 25 kb resolution, we discovered that genes in an active state exhibit characteristic median distance matrices, revealing distinctive higher-order genomic structures (Fig. 1e). To further elucidate the distinctions revealed in the median distance matrices, especially for genes in the active state, we applied t-Distributed Stochastic Neighbor Embedding (t-SNE) for dimensionality reduction. This allowed us to more effectively discern variations among genes with more than 10% active alleles within regions analyzed at a 25 kb resolution. The median distance matrices for the inactive state were similar to the overall matrices irrespective of transcriptional activity, whereas those in the active state diverged markedly (Fig. 1f). Furthermore, the distributions of median distance matrices in the active state demonstrated a tendency to segregate. This suggests that specific genes within a TAD can adopt distinct higher-order genomic structures when in an active state. These observations imply the formation of higher-order genomic structures that are specific to the active state. Each gene, when actively transcribed, displays distinctive higher-order genomic structures, suggesting that the influence of concurrent transcriptional activity with other genes on the same allele on these structures is probably very limited. To investigate this, we utilized seq-RNA-FISH to quantify the presence of transcriptional spots between genes. Our findings revealed that less than 10% of gene pairs exhibited concurrent transcriptional spot presence within the same allele (Supplementary Fig. 4). These results indicate that the higher-order genomic structures, which form in association with specific gene transcriptional states, are minimally influenced by the transcriptional states of adjacent genes (Supplementary Fig. 4).

### Relationship between cellular state and higher-order genomic structure

Higher-order genomic structure changes during cellular differentiation^19^. In mouse ES cells, a significant reduction in cell-to-cell heterogeneity in gene expression and upregulation of pluripotency marker genes, including *Nanog*, has been noted when cultured in medium containing MEK and GSK3β inhibitors (2i medium), as opposed to a serum/LIF medium^20^. To explore the relationship between these changes in cellular states and higher-order genomic structure surrounding the *Nanog* locus, we cultured mouse ES cells in 2i medium, conducted seq-DNA/RNA/IF-FISH analyses, and obtained data from 262 cells and 484 alleles (Fig. 2a-c). We performed t-SNE analysis on the median distance matrices of genes in their active or inactive states under both serum/LIF and 2i conditions. Strikingly, we found that the global median distance matrices in serum/LIF and 2i mediums clustered distinctly, indicating a low level of similarity (Fig. 2d). Moreover, similar to cells cultured in serum/LIF medium (Fig. 1f), those in 2i medium exhibited globally similar higher-order genomic structures in the inactive state of various genes, whereas characteristic structures were discernible in their active states. These observations strongly support the conclusion that the higher-order genomic structure varies depending on culture conditions and assumes distinct configurations specifically in the active states of genes.

**Fig. 2.**
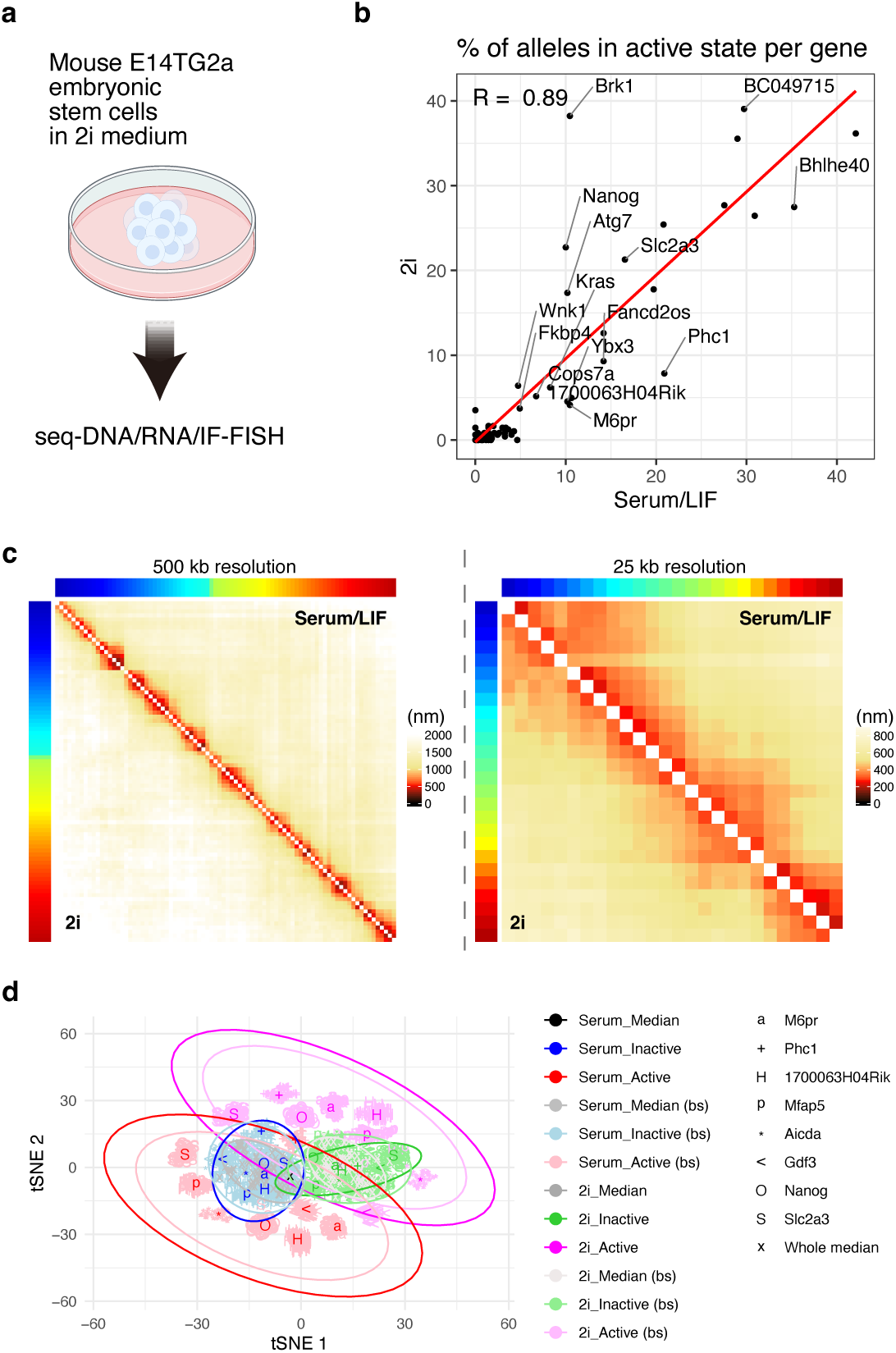
Relationship between cellular state and higher order genomic structure. **a** Schematic representation of mouse ES cells cultured in 2i medium and subjected to seq-DNA/RNA/IF-FISH analysis. **b** Proportion of cells exhibiting transcriptional bursting (burst frequency) in mouse ES cells cultured in serum/LIF and 2i media. Serum/LIF: *N* = 1,100; 2i: *N* = 484. Overall, high correlation values were observed, although the expression levels of certain genes varied substantially depending on the culture conditions. For instance, it is well-known that *Nanog* expression increases under 2i conditions; our study confirmed a corresponding increase in burst frequency in 2i culture. **c** Comparative analysis of higher-order genomic structures in mouse ES cells cultured in different media. Serum/LIF: *N* = 1,100; 2i: *N* = 484. No notable differences were observed at a 0.5 Mb resolution, whereas subtle structural variations were suggested at a 25 kb resolution. **d** t-SNE analysis was performed on the median distance matrices. Genes were categorized as either active or inactive, and their corresponding median distance matrices were utilized for the analysis. “Median” refers to the data when no active/inactive categorization was made. Using bootstrap methods (bs), we randomly sampled from the alleles, allowing for duplicates, and created median distance matrices the same number of times as the sample size *N*. The results from 100 bootstrap samples are shown.

### Formation of transcriptional regulatory factor condensates near genes in an active state

To elucidate the spatial relationship between the accumulation and localization of transcriptional regulatory factors/PTMs and transcriptional activity, we conducted a focused analysis of seq-IF-FISH data (Fig. 3a and Supplementary Fig. 5a). From 20 targets, we excluded those with low relative fluorescence intensities compared to negative controls, focusing our investigation on ten different proteins and PTMs (Supplementary Fig. 5a-c). At a 1 Mb resolution, we observed a positive correlation between the fluorescence intensities of transcriptional regulatory factors/PTMs at 3D coordinate of specific genomic loci, as determined from seq-DNA-FISH, and the ChIP-seq enrichment data (Supplementary Fig. 5d-g, see Methods). This strongly suggests that our methods can quantify the degree of protein accumulation in particular genomic loci.

**Fig. 3.**
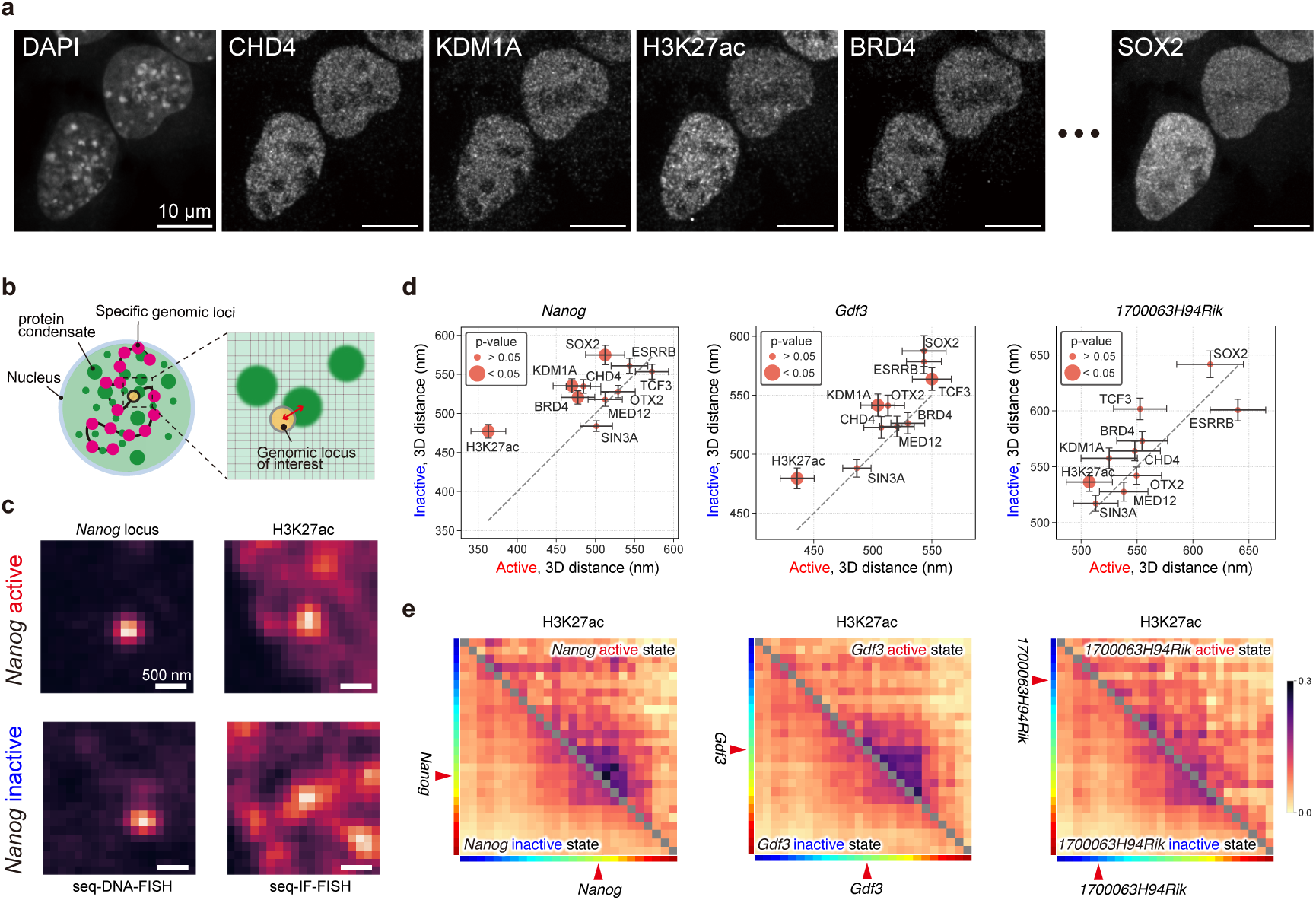
Formation of transcriptional regulatory factor condensates near genes in an active state. **a** Example images of nuclear localization of several proteins and post-translational modifications (PTMs) obtained by seq-IF-FISH analysis. The scale bars indicate 10 µm. For additional images, see Supplementary Fig. 5. **b** Schematic representation elucidating the spatial relationship between specific genomic regions and protein/PTM condensates. **c** Example images displaying the localization of H3K27ac obtained by seq-IF-FISH, in relation to the *Nanog* locus, obtained by seq-DNA-FISH. Transcription state of *Nanog* was determined by using seq-RNA-FISH data. **d** Median distances between specific genes and their closest condensates of transcriptional regulatory factors or PTMs, categorized by the transcriptional activity state of the genes. Error bars represent standard error. Sample sizes for each gene in active (inactive) states are as follows: *Nanog*: *N* = 110 (990); *Gdf3*: *N* = 217 (883); *1700063H04Rik*: *N* = 117 (983). **e** Colocalization probability matrix denoting the presence of H3K27ac condensates proximal to two distinct genomic loci pairs, with colocalization defined as being within 350 nm of each other. Red arrowheads indicate the location of the genes of interest.

We recognize the challenge in quantifying accumulation at resolutions below 1 Mb due to the diffraction limit. Thus, alternative analytical methods are essential. In previous live imaging studies, we showed that, in the active state of *Nanog*, BRD4 forms condensates in its spatial vicinity^11^. In this study, ’condensates’ refers to focal assemblies of identical proteins or PTMs, not necessarily indicating formation through liquid-liquid phase separation (LLPS) or similar processes. Using seq-IF-FISH data, we measured the distances from specific genes to their nearest condensates and found that, in the active state of *Nanog*, BRD4, histone H3 lysine 27 acetylation (H3K27ac), SOX2, and KDM1A were significantly concentrated in in the spatial vicinity of *Nanog*, forming condensates (Fig. 3b-d). In contrast, for genes *1700063H04Rik* and *Gdf3*, we found that H3K27ac alone and in combination with KDM1A and TCF3, respectively, formed spatially proximal condensates specifically in their active states (Fig. 3d). Further, we observed distinct differences between the active and inactive states in the colocalization probability of H3K27ac condensates with two distinct genomic loci pairs, defined as being within 350 nm of each other, especially in regions close to the target genes (Fig. 3e). These findings strongly support the notion that various factors form condensates in proximity to genes when they are active, and that these factors can vary from gene to gene^10^.

### Transcriptional regulation by dynamic interaction of distal enhancer-promoter

To elucidate how higher-order genomic structures and the formation of transcriptional regulatory factor/PTM condensates contribute to transcriptional activity, we focused on *Nanog*. Upon comparative analysis of median distance matrices according to the transcriptional activity states of the *Nanog*, we observed a notable difference in proximity between the upstream regions and the *Nanog* locus depending on the transcriptional state. Specifically, in the inactive state, the *Nanog* locus appeared to be spatially separated from the regions located more than 45 kb upstream from it, delineated by a known superenhancer referred to as −45SE^21^. In contrast, during the active state, we found increased proximity between the *Nanog* locus and regions upstream of −45SE (Fig. 4a). Contact probabilities (< 350 nm) revealed increased interaction frequencies with *Nanog*, extending up to the −190 kb region (Fig. 4b). When cumulative distance distribution between *Nanog* locus and other regions was examined, a statistically significant bias toward shorter distances was observed in the active state for the upstream −150 kb region. While not statistically significant, a similar trend was also apparent for the upstream −190 kb region, implying a functional role in *Nanog* transcriptional activity (Fig. 4c).

**Fig. 4.**
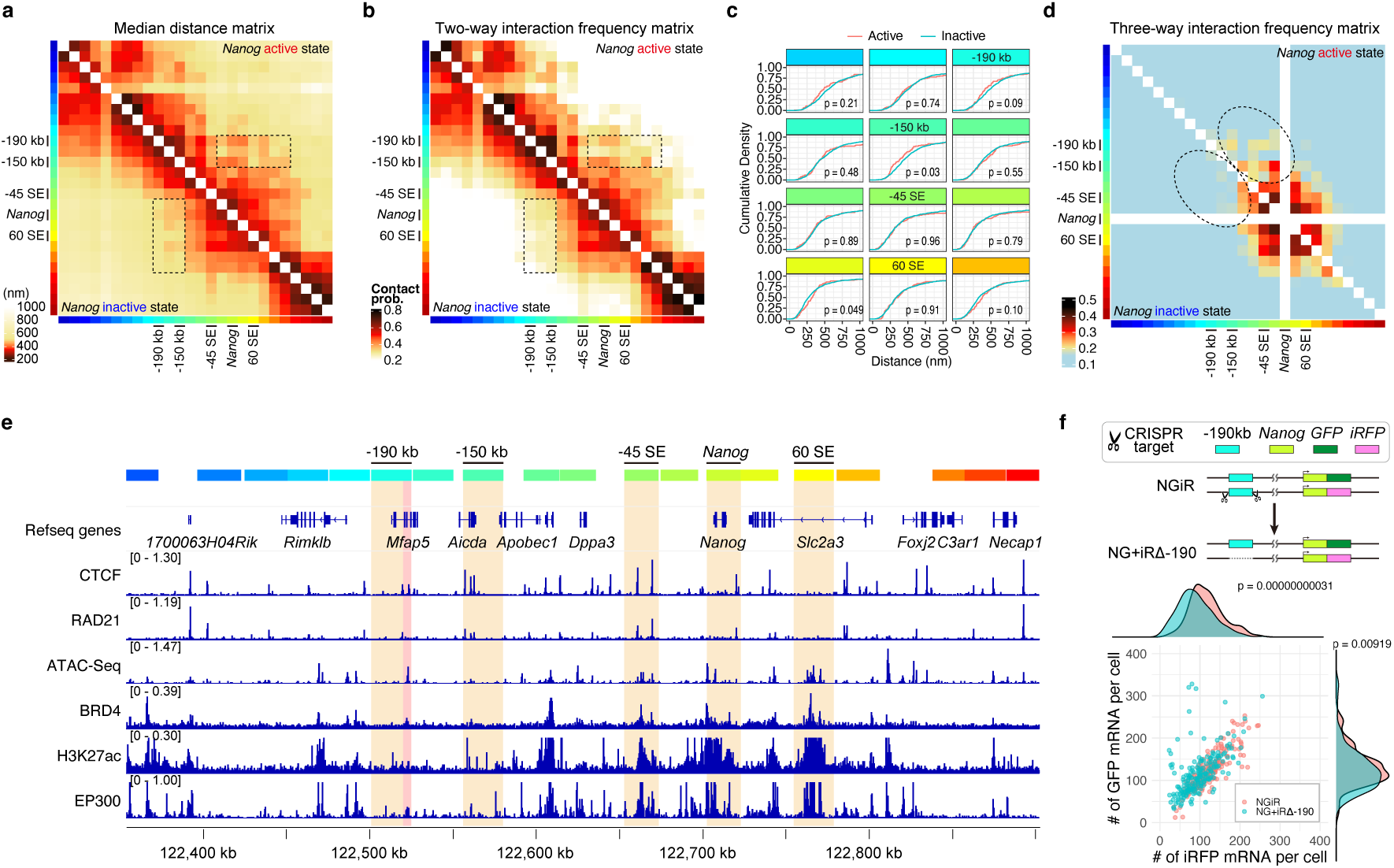
Transcriptional regulation by dynamic interaction of distal enhancer-promoter. **a** Median distance matrix for *Nanog* when in active or inactive transcriptional states. Sample sizes for active (inactive) states of *Nanog* are denoted as *N* = 110 (990). **b** Interaction frequency matrix for *Nanog* based on transcriptional activity states. The number of samples for *Nanog* in its active (inactive) states are as follows: *N* = 110 (990). Interactions were considered established when the distance between specified regions was 350 nm or less. **c** Cumulative density distribution of 3D distances between *Nanog* and specific genomic regions. Sample sizes for active (inactive) states of *Nanog* are indicated as *N* = 110 (990). *P*-values were determined using a two-sample Kolmogorov-Smirnov test. **d** Three-way interaction probability matrix between *Nanog* and two specific genomic regions. The proportion of instances where *Nanog* and the two other genomic regions are located within 350 nm of each other was calculated. **e** Epigenomic factor localization around the *Nanog* locus. Open chromatin domains characterized by ATAC-seq data, as well as BRD4 and EP300 localizations, are evident in the gene body region of *Mfap5*, indicating enhancer-like features. **f** Allele-specific single-molecule RNA-FISH analysis in NGiR and NG+iRΔ-190 cell lines. NGiR: *N* = 196; NG+iRΔ-190: *N* = 198. P-values were determined using a two-sided Wilcoxon rank sum test.

We further calculated the probability of simultaneous interactions (< 350 nm) between the *Nanog* locus and two other arbitrary genomic regions (Fig. 4d). Although we observed no substantial differences in the proximity near the *Nanog* region between active and inactive states, a marked increase was noted for combinations involving the upstream −150 and −190 kb regions. All these regions demonstrated increased three-way interactions with *Nanog* and known SEs like −45SE and another 60 kb downstream *Nanog* SE, 60SE^21^. This implies that the regions extending from −150 to −190 kb are specifically spatially proximal to the *Nanog* locus when it is in an active state. Notably, despite its genomic distance from *Nanog*, the −190 kb region exhibited elevated frequencies of three-way interactions. This region contains signals for open chromatin (as a proxy for ATAC-seq data), BRD4, and the histone acetyltransferase EP300 (based on ChIP-seq data), suggesting the presence of potential enhancer activity (Fig. 4e).

To investigate the functional role of the −190 kb region in *Nanog* transcription, we attempted its deletion via CRISPR genome editing (Fig. 4f and Supplementary Fig. 6). Using the NGiR cell line, where *GFP* and *iRFP* were knocked into the *Nanog* gene ^10^, we established a cell line with the −190 kb region deleted from the *Nanog-iRFP* allele, termed NG+iRΔ-190 (Supplementary Fig. 6). Quantification of transcriptional activity for each allele using single-molecule RNA FISH (smRNA-FISH) revealed a significant decrease in the number of *Nanog-iRFP* mRNA molecules originating from the allele lacking the −190 kb region (Fig. 4f). This suggests that the −190 kb region functions as one of multiple enhancers for *Nanog*, including the −45SE.

The formation of higher-order genomic structures is regulated by cohesin-mediated loop extrusion and its inhibition at binding sites by CTCF^22,23^. Considering the existence of multiple CTCF and cohesin binding sites around the *Nanog* region (Fig. 4e), we postulated that dynamic chromatin looping might serve as a regulatory mechanism for dynamic *Nanog* transcription^24,25^. Re-evaluation of publicly available Micro-C data following acute depletion of CTCF or RAD21, a cohesin subunit^26,27^, showed that acute CTCF depletion increased interaction frequencies between the *Nanog* region and multiple other regions, including −190 kb (Supplementary Fig. 7a,b). Conversely, acute depletion of RAD21 led to a substantial decline in such interactions (Supplementary Fig. 7c). Although care should be taken when interpreting these data, as they represent averages across multiple cells, our findings suggest that *Nanog* interactions are likely mediated by cohesin-driven loop extrusion. This mechanism seems to be inhibited near the −45SE region by CTCF (Supplementary Fig. 7a). Our data suggest that in the active state of *Nanog*, this inhibitory effect might be alleviated, thereby facilitating increased interactions with upstream regions.

### Dynamics of higher-order genomic structures and E-P communication

Alterations in higher-order genomic structure, which are dependent on transcriptional activity and include interactions between promoters and multiple distal enhancers, may impact the local dynamics of both promoters and enhancers. Our previous work has demonstrated that *Nanog* locus exhibits reduced mobility when in an active state^11,12^. Additionally, a growing body of evidence suggests that inhibiting transcription can enhance genomic dynamics, potentially stabilizing specific higher-order genomic configurations in a transcription-dependent manner^28,29^.

To investigate this further, we utilized a computational polymer modeling approach called PHi-C^30,31^. PHi-C enables us to convert input contact probability matrices obtained from seq-DNA/RNA-FISH data into the higher-order genomic structures and dynamics in both the transcriptionally active and inactive states of *Nanog* (Fig. 5a). Remarkably, the PHi-C structural modeling yielded data that exhibited strong correlations with seq-DNA-FISH-derived contact probabilities (Fig. 5a). While the PHi-C can theoretically predict the mean-square displacement (MSD) as a measure of mobility for specific genomic regions, the spatiotemporal scales are normalized by not pre-determined physical parameters such as the friction coefficient *γ* and contact distance *σ*. To recover the model in actual physical scales, we fitted the theoretical function of the PHi-C model with our experimental MSD data for the *Nanog* region^11^ under setting *σ* = 350 nm, thereby obtaining the value of *γ* kg/s (Fig. 5b). Astonishingly, the active state manifested a five-fold higher friction coefficient compared to the inactive state (*Nanog* active state: 3.08×10^-6^ kg/s; *Nanog* inactive state: 6.37×10^-7^ kg/s). This elevated friction coefficient implies not only a high level of viscosity but also suggests viscous interactions between the *Nanog* region and its surrounding environment (Fig. 5b and Supplementary Movies 2 and 3).

**Fig. 5.**
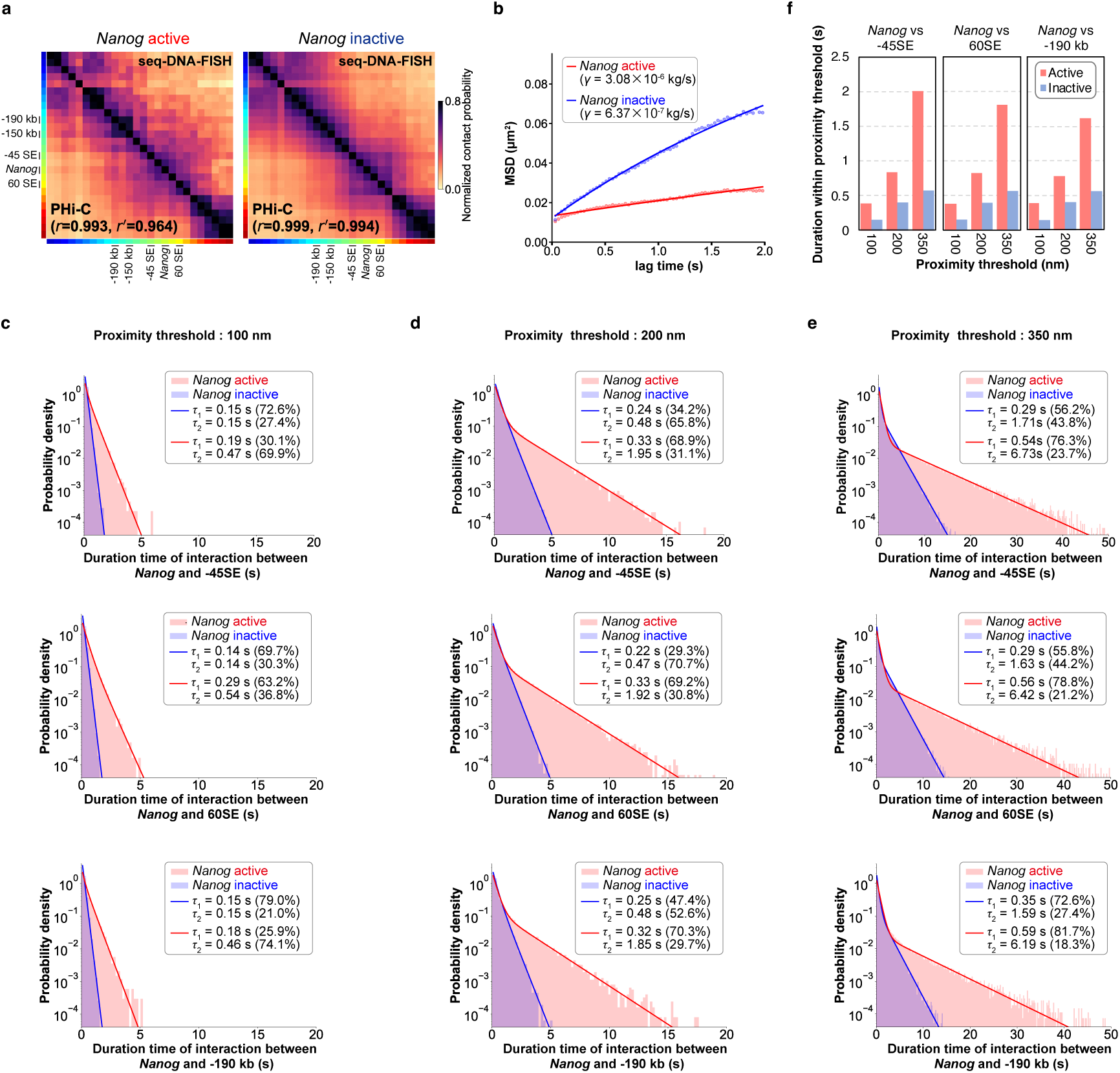
Dynamics of higher-order genomic structures and E-P communication. **a** Computational reconstruction of higher-order genome architecture utilizing PHi-C, informed by an interaction frequency matrix generated from seq-DNA/RNA-FISH data. The top right panel represents the seq-DNA-FISH data, while the bottom left showcases the interaction frequency matrix as estimated through PHi-C. Interaction frequency matrices are distinctly displayed, based on *Nanog*’s active and inactive states. **b** Mean-square displacement (MSD) of the *Nanog* locus. MSD values from PHi-C were fitted to the experimental data for the *Nanog* region^11^ to determine the friction coefficient *γ*. Further details can be found in Methods. **c** Distribution of the duration of interaction between *Nanog* and −45SE, between *Nanog* and 60SE, and between *Nanog* and −190 kb at a proximity threshold of 100 nm. **d** Distribution of the duration of interaction between *Nanog* and −45SE, between *Nanog* and 60SE, and between *Nanog* and −190 kb at a proximity threshold of 200 nm. **e** Distribution of the duration of interaction between *Nanog* and −45SE, between *Nanog* and 60SE, and between *Nanog* and −190 kb at a proximity threshold of 350 nm. **f** Duration of closeness below the proximity threshold between *Nanog* and −45SE, between *Nanog* and 60SE, and between *Nanog* and −190 kb.

Consistent with the transcription hub model, our data provide evidence that various factors form condensates near active genes (Fig. 3). This fivefold increase in the friction coefficient suggests that condensate formation influences the dynamics of not only the gene in the active state but also other genomic loci (Supplementary Movies 2 and 3)^32^. Recent studies employing Micro-C have revealed characteristic interactions between the +1 nucleosome downstream of the transcription start sites of enhancers and promoters^8^. These nucleosomes are inferred to be in remarkably close proximity (< 100s nm), suggesting that such states are maintained for a certain extent of time. To elucidate the proximity dynamics between the *Nanog* locus and −45SE, we measured the duration of closeness, using 100, 200, and 350 nm as proximity thresholds in the simulation. Remarkably, at each of these thresholds, we found that the duration of interactions within the threshold distance was more than doubled in the active state compared to the inactive state (Fig. 5c-f). Notably, not only was this true for −45SE, but the durations also increased for other surrounding enhancer regions like −190 kb and 60SE (Fig. 5c-f). Consequently, it can be posited that in the active state of *Nanog*, sustained contacts with multiple enhancer regions induce continuous transcriptional activity.

We had also observed that the dynamics of *Sox2* are less mobile in the active state^11^. Re-analysis of chromatin tracing data for the *Sox2* region^33^ revealed the contact duration with a downstream SE located 100 kb downstream more than doubled during the active state (Fig. 6).

**Fig. 6.**
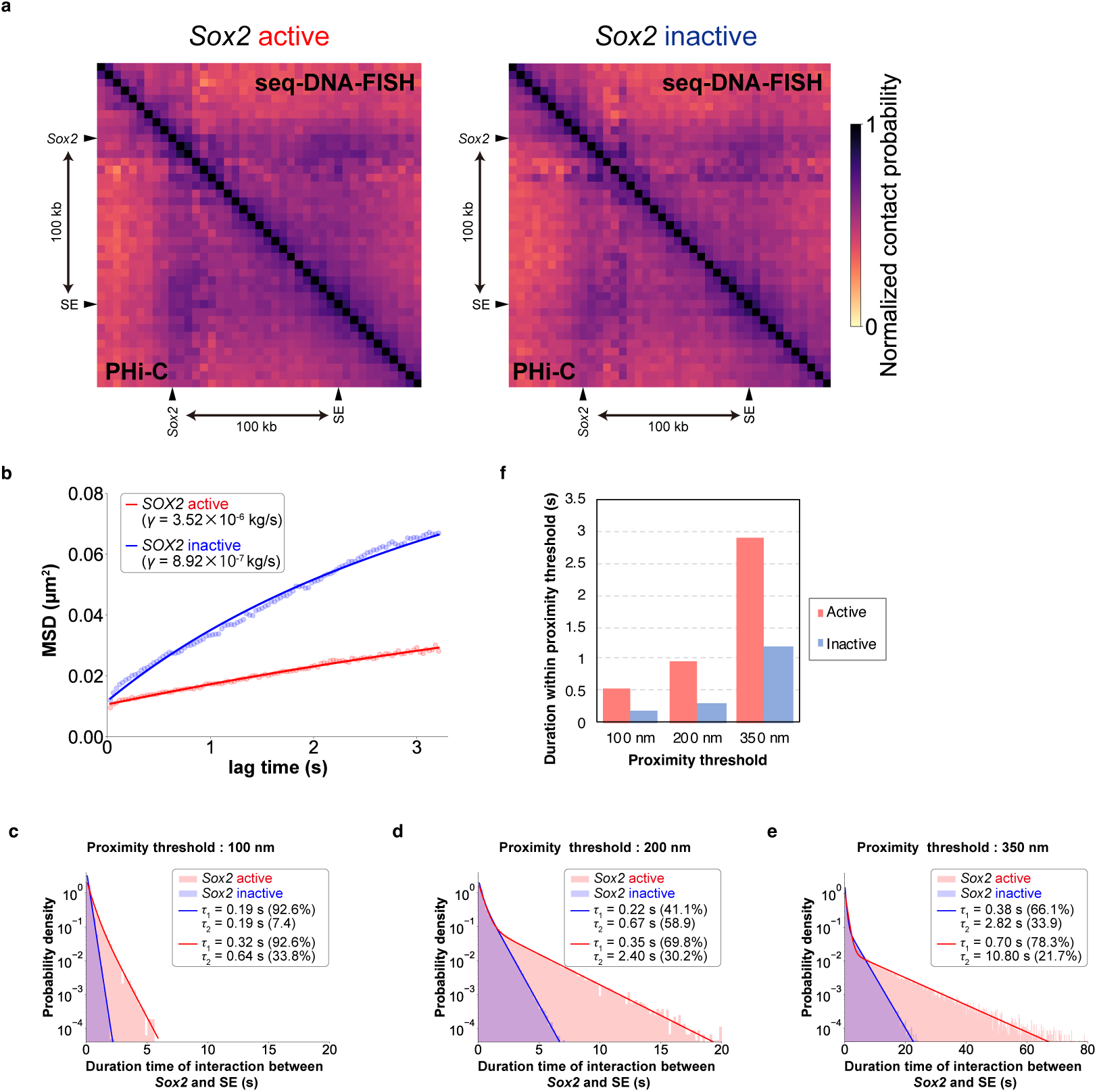
Estimation of higher-order genomic structural dynamics for each *Sox2* transcriptional activity state by PHi-C. **a** Interaction frequency matrix derived from chromatin tracing data^33^ for *Sox2* transcriptional activity, with a threshold set at 350 nm, used to estimate higher-order genome architecture through PHi-C. The top right panel shows the seq-DNA-FISH data, and the bottom left depicts the interaction frequency matrix as estimated by PHi-C. **b** Mean-square displacement (MSD) of the *Sox2* locus. MSD values from PHi-C were fitted to the experimental data for the *Sox2* region^11^ to determine the friction coefficient *γ*. Further details can be found in Methods. **c** Distribution of the duration of interaction between *Sox2* and SE at a proximity threshold of 100 nm. **d** Distribution of the duration of interaction between *Sox2* and SE at a proximity threshold of 200 nm. **e** Distribution of the duration of interaction between *Sox2* and SE at a proximity threshold of 350 nm. **f** Duration of closeness below the proximity threshold between *Sox2* and SE.

In summary, our research compellingly demonstrates that within specific gene regions, alterations in higher-order genomic architecture coupled with an increase in the local viscosity due to the formation of transcription hubs serve to prolong the duration of E-P interactions in the active state. This paradigm-shifting insight offers a nuanced understanding of the mechanistic intricacies governing transcriptional regulation (Fig. 7).

**Fig. 7.**
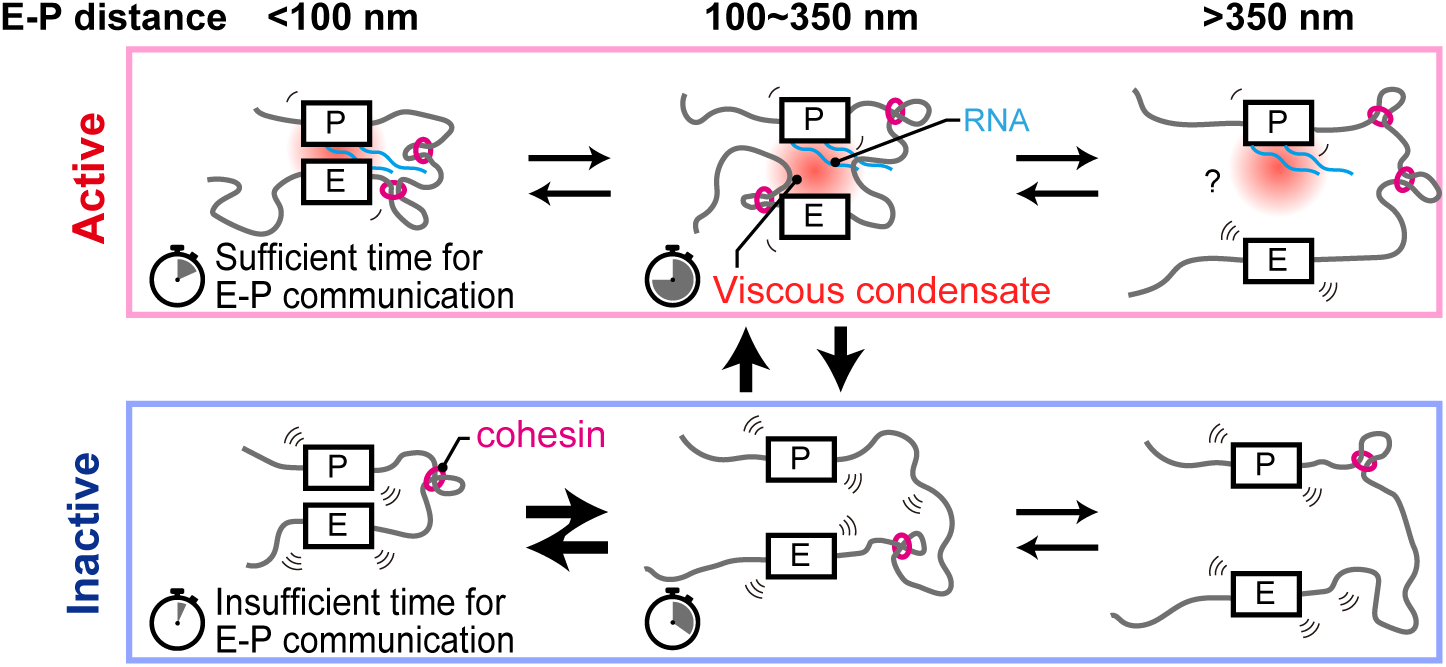
Transcriptional model incorporating increased E-P interaction durations specific to the active state. In the active state, the specific higher-order genomic structures are formed, along with the assembly of condensates mediated by proteins and RNAs. This results in an elevated-viscosity environment around the gene, extending the duration of E-P interactions. In the inactive state, viscosity is lower, leading to shorter E-P interactions compared to the active state. In both transcriptional states, the dynamics of higher-order genomic structures are facilitated by cohesin-mediated loop extrusion. Timer icons represent the average duration of each state.

## Discussion

In the present study, we applied sequential DNA/RNA/IF-FISH techniques to elucidate the higher-order genomic structures and local transcriptional regulatory factor condensates in the active state of specific genes. Computational simulations compellingly supported our empirical findings, demonstrating that these higher-order genomic structures elevate local viscosity. This, in turn, extends the duration of E-P interactions (Fig. 7).

Our results find additional support in the model proposed by Zuin et al., who accounted for the non-linear relationship between E-P interaction frequencies and RNA expression levels through E-P communication-mediated transcriptional bursting^34^. Notably, our findings, with 100 nm and 200 nm as thresholds, showed average proximity durations of approximately 0.4 and 0.8 seconds, aligning well with the E-P interaction timescales described by Zuin et al.^34^. Despite this consistency, we observed significant variability in higher-order genomic structures even in the active state (Fig. 4c). This suggests that several factors, such as the gene region’s higher-order genomic structure, E-P genomic distance, and promoter activity strength, contribute to the lack of a straightforward correlation between transcriptional activation and E-P physical distance^2–6^ (Fig. 7). Building on these observations, our findings lend principal support to a transcriptional hub model^7^. In our model, changes in higher-order genomic structure and viscosity lead to the formation of hubs approximately 350 nm in size. Our model also accommodates recent observations that reported E-P interactions at distances of less than hundreds of nanometers^8^ (Fig. 7).

We speculate that extended interaction times facilitate E-P communication, which has several implications for the recruitment of transcription factors and coactivators, as well as the assembly of Mediator complexes^35^. Intriguingly, our data suggest that E-P contact frequency alone may not suffice for understanding functional relationships with transcriptional activity (Figs. 5, 6), as we observed substantial changes in E-P contact duration between active and inactive states (Figs. 5f, 6f).

However, our study is not without limitations. We were unable to experimentally track temporal changes in the same cell and allele, suggesting that static methodologies like Hi-C and FISH may not capture all relevant dynamics. These constraints underline the necessity for developing sophisticated imaging techniques that enable multicolor, high-temporal-resolution (< 50 ms intervals) imaging.

While the initiation mechanisms for condensate formation remain to be clarified, there is a possibility that condensates not only contain transcriptional regulatory factors but also nascent RNA and eRNAs transcribed from enhancers. Recent reports have shown that RNA-containing condensates contribute to the rise in local viscosity^32^. This suggests the potential role of nascent RNA and eRNA as crucial elements in higher-order genomic structuring and E-P interactions, paving the path for future research.

## Methods

### Cell lines

The E14tg2a cell line (AES0135, Riken Cell Bank, JP) was cultured at 37℃ and 5% CO_2_, on either laminin-511 (LN511) (BioLamina, Stockholm, Sweden) or a gelatin-coated dish. Two types of mediums were used: serum/LIF medium, which included Dulbecco’s modified Eagle’s medium (DMEM, Wako, Osaka, Japan, 197-16275), 15% fetal bovine serum (FBS, GE Healthcare, Little Chalfont, UK, SH30396.03), 0.5 mM monothioglycerol solution (Wako, 195-15791), 1×MEM nonessential amino acids (Wako, 139-15651), 2 mM L-alanyl-L-glutamine solution (Wako, 016-21841), 1,000 U/mL leukemia inhibitory factor (Wako, 195-16053), and 20 µg/mL gentamicin (Wako, 078-06061); and 2i medium, which contained the same ingredients as the serum/LIF medium along with 3 µM CHIR99021 (Cayman Chemical, Ann Arbor, MI, USA, 13122) and 1 µM PD0325901 (Chemscene, Monmouth Junction, NJ, USA, CS-0062).

C57BL/6J mESCs (Bruce 4 C57BL/6J, male, EMD Millipore, Billerica, MA) and other knock-in (KI) derivatives, including the NGiR cell line (160329-B6-2 GFP/iRFP+4)^10^ and NG+iRΔ-190 cell line, were cultured under the same conditions, using either LN511 or a gelatin-coated dish and either serum/LIF or 2i medium.

### Plasmids

In the present study, plasmids were engineered employing conventional molecular biology techniques, and we utilized the following three plasmid variants. The sequences of these plasmids can be accessed through the respective URLs provided. For CRISPR-mediated gene editing, two CRISPR vectors were used: eSpCas9-EF-49_10_L1 (available at https://benchling.com/s/seq-nS1bLoWAlYFDGQ2DtaiK?m=slm-pCIkvw6Miu0Vhto7abmZ) and eSpCas9-EF-49_10_R1 (accessible at https://benchling.com/s/seq-McPdsNwm54kANzquOwyv?m=slm-7jSZnlfEwwcawOARrJ7e). Additionally, the mTagBFP2 expression vector, pCAG-mTagBFP2, was used (Addgene plasmid #122373; available at https://www.addgene.org/122373/). pCAG-mTagBFP2 can be sourced from Addgene (Watertown, MA, USA).

### Microscopy

Images were acquired using a Nikon Ti-2 microscope with a CSU-W1 confocal unit (Yokogawa), a 100× Nikon Plan Apo λ oil-immersion objective lens (NA 1.4), and an iXon Ultra EMCCD (Andor Technology), operated using NIS-Elements software (ver. 5.11.01; Nikon). The microscope was also equipped with 405, 488, 561, and 637 nm lasers (Andor Technology), and an ASI MS-2000 piezo stage (ASI). Z-stack images spanning 10 µm with 200 nm intervals (51 sections; 130 nm/pixel) were acquired.

### Validation of fluorescent detection method in sequential RNA-FISH

C57BL/6J murine embryonic stem cells (mESCs) were seeded at a density of 7.5 × 10^4^ cells in four wells of an 8-well chambered cover glass with #1.5 glass, coated with LN511 (Cellvis, Sunnyvale, CA, USA, C8-1.5HN). Cells were fixed the following day with 4% paraformaldehyde (PFA) for a 10-minute incubation. This was followed by three washes with 1× Phosphate-Buffered Saline without Calcium and Magnesium (PBS), a permeabilization step for one hour at −20°C in 70% ethanol, and air-drying for 10 minutes. Subsequent treatment with 0.5% Triton-X in PBS was performed at room temperature for 15 minutes, after which the cells were washed three times with PBS. Cells were then blocked at room temperature for 15 minutes in a solution containing PBS, 10 mg/ml UltraPure BSA (Invitrogen AM2616), 0.3% Triton-X, 0.1% dextran sulfate (Nacalai Tesque, Kyoto, Japan, 03879-72), and 0.5 mg/ml sheared Salmon Sperm DNA (Invitrogen AM9680). Following blocking, samples were washed with 2×SSC (Nacalai Tesque, Kyoto, Japan, 32146-91).

Hybridization was performed at 37°C for 12 hours in a humidified chamber using seq-RNA-FISH probe sets (see below) and 10 nM polyT LNA oligonucleotide (Supplementary Table 1) in 50% hybridization buffer. This buffer consisted of 50% formamide (Wako, 066-02301), 2×SSC, and 10% (w/v) dextran sulfate. Post-hybridization, samples were washed at room temperature for 30 minutes with a 55% wash buffer, followed by three rinses with 4×SSC.

Hybridization with either 100 nM readout probe or a mixture of secondary probe and readout probe was executed in 10% EC buffer (10% ethylene carbonate (Sigma-Aldrich, E26258), 10% dextran sulfate (Sigma-Aldrich, D4911) and 4×SSC) at room temperature for 20 minutes. The samples were then washed with 4×SSCT (4×SSC and 0.1% Triton-X) and subsequently with 12.5% wash buffer (12.5% formamide, 2× SSC and 0.1% Triton X-100) at room temperature for 30 seconds. Additional staining with DAPI solution (4×SSCT solution with DAPI (1:100, Dojindo, 340-07971)) was carried out for 30 seconds at room temperature. Samples were imaged under a microscope after the addition of an anti-bleaching buffer (50 mM Tris-HCl pH 8.0, 300 mM NaCl, 2xSSC, 3 mM trolox (Sigma-Aldrich, 238813), 0.8% D-(+)-Glucose (Nacalai Tesque, Kyoto, Japan, 16806-25),1,000-fold diluted catalase (Sigma-Aldrich, C3155), 0.5 mg ml−1 glucose oxidase (Sigma-Aldrich, G2133)). Post-imaging, samples were treated for two minutes at room temperature with 55% wash buffer (55% formamide, 2× SSC and 0.1% Triton X-100) to facilitate stripping, washed twice with 4×SSCT, and reprocessed from hybridization to imaging. Post-stripping verification was confirmed by imaging the samples treated with DAPI solution and anti-bleaching buffer. Probe sequences utilized in this study are listed in Supplementary Table 2.

### Readout and secondary probe design and synthesis

In this study, we generated one million 15-base probes using the numpy.random function in Python. From this pool, we selectively isolated those with 5’ ends starting with either adenine (A) or thymine (T) and a GC content within the range of 40-60%. Indices were crafted from human and mouse transcriptome data available from NCBI. The generated probes were then mapped to these indices in sequential fashion—first to the human and then to the mouse transcriptome—utilizing Bowtie2 in its default mode (search for multiple alignments, report the best one). Probes failing to map were retained for further analysis. From these, a subset of 5,000 was chosen and subjected to BLAST analysis via its web interface, sequentially against the human and mouse transcriptomes to eliminate those demonstrating high complementarity. Further refinement was carried out using blastn in BLAST+ to exclude probes with high complementarity (those matching 10 bases or more). Of the remaining probes, 227 were employed as secondary probes and/or readout probes. Each secondary probe was designed to include sequences complementary to the primary probe, in addition to two identical readout probe binding sequences. The secondary probes were custom synthesized and purified using an OPC column by Eurofins. Readout probes tagged with Alexa 488 and Alexa 647 at their 5’ ends were synthesized by Thermo Scientific, while those labeled with ATTO565 at the 5’ end was synthesized by Sigma Aldrich. All readout probes underwent purification through High-Performance Liquid Chromatography (HPLC).

### Primary probe design for sequential DNA/RNA-FISH

Target regions were selected based on their interactions with *Nanog* in mouse embryonic stem (ES) cells. This was guided by evidence from 4C-seq data^15^(Supplementary Fig. 2a) as well as promoter-capture Hi-C analyses in mouse ES cells^36^. Aside from the aforementioned *Nanog*-interactive regions, other regions of interest were set at approximately 0.5 Mb intervals to comprehensively cover the genome landscape. Furthermore, we also designated 28 regions at intervals of 25 kb. All target regions utilized for seq-DNA-FISH were cataloged in Supplementary Table 3.

Probe sequences were generated by selecting 35-nucleotide sections from target regions, focusing specifically on GC content that lies outside the 45–65% range. For the design of primary probes, the target genomic region was obtained from the unmasked and repeat-masked GRCm38/mm10 mouse genome FASTA files, downloaded from Ensembl release 102. Probe design was facilitated through Oligominer (https://github.com/beliveau-lab/OligoMiner)^37^, employing parameters set as “-l 35 -L 35 -g 45 -G 65 -t 37 -s 300 -S 5 -c 5 -C 1”. Subsequently, Bowtie2 was utilized to identify similar sequences with parameters specified as “-t -k 11 --local -D 20 -R 3 -N 0 -L 19 -i C,1 -S”. Following the initial probe design, alignment to the unmasked mouse genome was performed using Bowtie2 for off-target evaluation. Criteria for off-target hits were specified as any alignment with a minimum of 19 matched bases that resided outside the genomic coordinates of the designated target region. Probes yielding over 10 off-target matches were excluded. Concurrently, probes were scrutinized against a BLAST database formulated from common repeating sequences in mammals. The FASTA file for “Simple Repeat” sequences restricted to “Mammalia only” was sourced from Repbase^38^. Probes with at least 19 matched bases to the repeats index were disregarded.

Subsequent to this, cross-hybridization candidates were eliminated using blastn. Probes pairs displaying at least 19 matched bases were discarded in the final probe selection process. The finalized probe sets were meticulously chosen to preserve probe specificity and to ensure a relatively uniform distribution of probes along the target sequence. Probes located near the central region of the target genomic region received preferential selection. For each genomic locus targeted, a range of 100 to 200 primary probes were employed within a single target genomic region to visualize individual loci as diffraction-limited spots, as per seq-DNA-FISH methodology.

The primary probes for seq-DNA-FISH are composed of 35-nucleotide (nt) sequences specific to the genomic region, accompanied by four identical 15-nt secondary probe binding sites. Additionally, these probes possess 20-nt primer binding sites situated at both the 5’ and 3’ termini. The 15-nt secondary probe binding sites are associated with one of the 40 sequential rounds utilized for diffraction-limited spot imaging (Fig. 1a). The target sequences for these probes are listed in Supplementary Table 4.

As fiducial markers, primary probe targeting the repetitive sequence located in the *3632454L22Rik* locus on the X chromosome were utilized. This fiducial probe is similarly constructed, containing 35-nt genomic-specific sequences, albeit flanked by three distinct 15-nt secondary probe binding sites. These probes also possess a pair of 20-nt primer binding sites, identical to the previously described primary probes. The target sequence for the fiducial probe is AAGGAAGCCAGCTGTGGGTAAGGAAGCCAGCTGTG. Phosphorylated 5’ termini of this fiducial probe was synthesized by Eurofins and purified via HPLC.

For seq-RNA-FISH primary probe design, target candidates were initially selected from among the regions previously defined for seq-DNA-FISH, specifically focusing on those that included a transcription start site. Further refinement was based on gene expression levels; only those regions exhibiting non-negligible expression were included in the analysis. Expression levels were determined based on CEL-seq2 data from serum/LIF cultured mouse ES cells^10^, accessed at https://www.ncbi.nlm.nih.gov/geo/query/acc.cgi?acc=GSE132591, considering genes with a unique molecular identifier (UMI) count of 10 or more as suitable candidates for seq-RNA-FISH analysis. The target sequences were selected referencing PaintSHOP (https://oligo.shinyapps.io/paintshop/)^39^. Each targeted RNA was imaged with up to 48 primary probes per target RNA to visualize individual RNAs as diffraction-limited spots. The number of probes could vary for shorter genes, with the minimum being 15. These RNA-specific primary probes range from 30 to 37-nt in length and are flanked by three identical 15-nt secondary probe binding sites. They also contain 20-nt primer binding sites at both the 5’ and 3’ ends. The 15-nt secondary probe binding sites are associated with one of the 27 sequential rounds for diffraction-limited spot imaging. The targeted sequences for these seq-RNA-FISH probes are listed in Supplementary Table 5.

### Primary probe preparation for seq-DNA/RNA-FISH

Primary probes designed for seq-DNA/RNA-FISH, as described in the “Primary probe design for sequential DNA/RNA-FISH” section, were obtained as oligoarray complex pools from Twist Bioscience. These oligo pools served as templates and were amplified in accordance with previously reported methodologies^13^. For seq-DNA-FISH primary probes, amplification from the oligo pool was conducted using seqDNA1F and seqDNA1R primers, employing KOD One^®^ PCR Master Mix (TOYOBO, Japan). The resultant DNA was then purified using DNA Clean & Concentrator (Zymo Research). A second round of PCR amplification was executed using seqDNA2F and seqDNA2R primers, followed by in vitro transcription using the MEGAshortscript Kit (Thermo Fisher AM1354). Reverse transcription was facilitated using seqDNA_revT primer and Thermo Scientific Maxima H Minus Reverse Transcriptase (Thermo Fisher EP0751). After reverse transcription, the single-stranded DNA (ssDNA) probes were subjected to alkaline hydrolysis with 1 M NaOH at 65℃ for 15 minutes to degrade the RNA templates, subsequently neutralized with 1 M acetic acid, and then ethanol precipitated. These amplified primary probes were resuspended in seq-DNA-FISH primary hybridization buffer, which consisted of approximately 1 nM per probe, 100 nM 3632454L22Rik fiducial marker probe, 40% formamide, 2× SSC, and 10% (w/v) dextran sulfate. If any precipitation was observed, it was dissolved by heat treatment at 65℃ for 15 minutes. The probes were stored at −20℃ until use.

For seq-RNA-FISH primary probes, a similar amplification protocol was followed. However, the initial amplification from the oligo pool utilized seqRNA1F and seqRNA1R primers, while the second round of PCR employed seqRNA2F and seqRNA2R primers. For reverse transcription, seqRNA1F primer was used. All other procedures were analogous to those employed for the seq-DNA-FISH primary probes. The oligonucleotides utilized are listed in Supplementary Table 1.

### DNA–antibody conjugation

Genomic regions interacting specifically with the Nanog promoter region in mouse ES cells were identified as subjects for investigation, analyzed using Enrichment Analysis in ChIP-Atlas ^40^, based on findings previously published by de Wit et al. ^15^. Targets were selected based on low p-values and the availability of commercial antibodies not containing bovine serum albumin or gelatin. A list of targeted antigens and the corresponding antibodies used in this study is provided in Supplementary Table 6.

The preparation of oligonucleotide DNA-conjugated primary antibodies was conducted as previously described^41^. Briefly, antibodies (100 μg) underwent buffer exchange to PBS via 50-KDa Amicon Ultra Centrifugal Filters (Millipore, UFC505096) and were treated with 10 equivalents of PEGylated SMCC cross-linker (SM(PEG)2) (Thermo Scientific 22102) diluted in anhydrous DMF (Vector Laboratories S4001005). Following incubation at 4℃ for 2 hours, the solution was purified using 7K MWCO Zeba Spin Desalting Columns. Concurrently, 300 μM 5’ thiol-modified 18-nt DNA oligonucleotides—comprised of an AAA base sequence and a 15-base secondary probe binding sequence (Eurofins)—were reduced with 50 mM dithiothreitol in PBS at room temperature for 2 hours and purified using NAP5 columns (GE Healthcare 17-0853-01). Maleimide-activated antibodies were then combined with 11 equivalents of the reduced form of the thiol-modified DNA oligonucleotides in PBS and incubated at 4℃ overnight. The resulting DNA-primary antibody conjugates were washed four times with PBS and concentrated via 50-KDa Amicon Ultra Centrifugal Filters. The concentrations of the conjugated oligonucleotide DNA and antibodies were quantified using a BCA Protein Assay Kit (Thermo Scientific 23225) and Nanodrop (Thermo Scientific).

Functional validation of the conjugated antibodies was confirmed through SDS-PAGE and Coomassie Brilliant Blue (CBB) staining, which demonstrated the expected reduction in electrophoretic mobility due to oligonucleotide conjugation. Additional verification was conducted through immunofluorescence, utilizing fluorophore-conjugated secondary antibodies that recognize the host species of each antibody; successful nuclear staining affirmed the functionality of the oligonucleotide-labeled antibodies.

To achieve vendor-recommended concentrations for application, a mixture of 20 distinct antibodies was combined in a 1.5 ml tube. Subsequently, this mixture was brought to a final volume of 500 µL using PBS. DNA-primary antibodies were buffer-exchanged to PBS using 50-KDa Amicon Ultra Centrifugal Filters. Concentrations were verified using a BCA Protein Assay Kit and Nanodrop. These preparations were stored at −80℃ until use.

### seq-DNA/RNA/IF-FISH

Glass-bottom six-well plates (Cellvis, P06-1.5H-N) were equipped with cell culture inserts (ibidi, ib80209). Only the area within the cell culture inserts was treated with poly-d-lysine (PDL) (Sigma-Aldrich, P6407) at a concentration of 0.1 mg/mL for 60 minutes. Following the treatment, the plates were air-dried overnight. The inserts were subsequently coated with LN511 at 37°C for 1 hour. E14tg2a ES cells, pre-cultivated in either serum/LIF or 2i media, were seeded at a density of 2 × 10^4^ cells per insert and incubated at 37°C with 5% CO_2_. The next day, cells were fixed with 4% PFA for 10 minutes, followed by three washes with PBS. Supernatants were discarded, and cells were treated with 70% ethanol at −20°C for 1 hour, and air-dried for 10 minutes. Custom HybriWell Sealing Systems (Grace Bio Labs, Custom HBW13 FL_1, 13mm Diameter × 0.25mm Depth ID, 25mm x 28mm OD, 1.5mm Ports, Adhesive A12) were applied at this stage.

Cell permeabilization was initiated with a 15-minute treatment of 0.5% Triton-X in PBS at room temperature, followed by three PBS washes. Blocking was carried out for 15 minutes at room temperature using a blocking solution that included 10 mg/mL UltraPure BSA (Invitrogen AM2616), 0.3% Triton-X, 0.1% dextran sulfate (Nacalai Tesque, 03879-72), and 0.5 mg/mL sheared Salmon Sperm DNA (Invitrogen AM9680). DNA oligonucleotide-conjugated primary antibodies (Supplementary Table 6) were incubated in this blocking solution, with 100-fold diluted SUPERase In RNase Inhibitor (Invitrogen AM2694), at 4°C overnight.

Subsequent to three 10-minute washes with PBS, the samples underwent post-fixation with freshly prepared 4% PFA in PBS for 5 minutes at room temperature. This was followed by six washes with PBS and an additional post-fixation step using 1.5 mM BS(PEG)5 (Thermo Scientific, A35396) in PBS for 20 minutes at room temperature, quenched by 100 mM Tris-HCl pH 7.5 for 5 minutes. After washing with PBS, the samples were rinsed with 2×SSC (Nacalai Tesque, 32146-91). For hybridization, seq-RNA-FISH probe sets (1 nM each per probe) and 10 nM polyT LNA oligonucleotide (Supplementary Table 1) were combined in a 50% hybridization buffer consisting of 50% formamide (Wako, 066-02301), 2×SSC, and 10% (w/v) dextran sulfate. The hybridization process was carried out at 37°C for 48 hours in a humidified chamber. To prevent dehydration, the port seals accompanying the HybriWell™ Sealing System were applied. Following hybridization, samples were rinsed with a 55% wash buffer at room temperature for 30 minutes, then washed three times with 4×SSC. For imaging, please refer to the “seq-DNA/RNA/IF-FISH imaging” section.

Following seq-RNA-FISH imaging, the specimens underwent a preparation process for seq-DNA-FISH primary probe hybridization. Initially, the specimens were washed using PBS, and then subjected to a one-hour incubation at 37℃ in RNase A/T1 Mix (Thermo Fisher EN0551), diluted 100 times. Subsequent to this step, the samples were subjected to three rinses in PBS, and an additional three washes in a 50% denaturation buffer containing 50% formamide and 2×SSC; this was followed by a 15-minute incubation at room temperature. The samples were then exposed to a 90℃ heat treatment for 4.5 minutes in the 50% denaturation buffer, while sealing the inlet and outlet of the custom chamber with accompanying seals from the HybriWell™ Sealing System.

Post-heating, the samples were again rinsed with 2×SSC. The seq-DNA-FISH primary hybridization buffer, comprising approximately 1 nM per probe, a 100 nM 3632454L22Rik fiducial marker probe, 40% formamide, 2×SSC, and 10% (w/v) dextran sulfate, was applied and incubated at 37℃ for 72 hours within a humidified chamber. To prevent dehydration, the HybriWell™ Sealing System’s holes were sealed with the provided port seals. Subsequent to the hybridization, the samples were washed with a 40% wash buffer containing 40% formamide, 2×SSC, and 0.1% Triton X-100, at room temperature for 15 minutes. This was followed by three additional washes in 4×SSC. Thereafter, the samples underwent further processing to ’padlock’ the primary probes. For this purpose, a 31-nucleotide global ligation bridge (final concentration: 100 nM; Supplementary Table 1) was hybridized in a 20% hybridization buffer, composed of 20% formamide, 10% dextran sulfate (Sigma-Aldrich, D4911), and 4×SSC, at 37℃ for 2 hours. Following this, the samples were washed three times for a total of 15 minutes with a 10% wash buffer, consisting of 10% formamide, 2×SSC, and 0.1% Triton X-100. The specimens were then incubated with a 20-fold diluted Quick Ligase in 1x Quick Ligase Reaction Buffer from Quick Ligation Kit (New England Biolabs, M2200), supplemented with an additional 1 mM ATP (TAKARA Bio, 4041), at room temperature for 1 hour. This step was designed to facilitate the ligation reaction between the 5’- and 3’-ends of the seq-DNA-FISH primary probes. The samples were then subjected to a series of washes and incubations, including fixation and amine modification steps, to further stabilize the primary probes. Imaging for seq-DNAFISH and seq-IF-FISH was carried out as described in the subsequent section (see ‘seq-DNA/RNA/IF-FISH imaging’).

### seq-DNA/RNA/IF-FISH imaging

The sequential imaging procedure was carried out based on the methodology previously reported by Takei et al.^13^. The fluidics delivery system used for this process also adhered to prior research^18^. Briefly, an automated fluidics delivery system was constructed, comprising two multichannel fluidics valves (EZ1213-820-4; IDEX Health & Science) and a Hamilton syringe pump (63133-01, Hamilton Company). Integration of the fluidics valves and syringe pump with homemade connectors, as well as the coordination with microscope imaging, was managed via a custom Python script. Samples were mounted on the microscope, and tubes were positioned to allow the flow of solutions and waste fluids into the apertures of HybriWell seals. A volume of 500μL DAPI solution was flowed through the system to stain cell nuclei, followed by the capture of 30 to 50 fields of view (FOVs). Subsequently, a 400μL mixture of 100 nM secondary and readout probes was flowed through in 10% EC buffer (containing ethylene carbonate, dextran sulfate, and 4×SSC) and hybridized at room temperature for 20 minutes. This was followed by a brief room temperature wash, using 500 μL of 4×SSCT (4× SSC and 0.1% Triton-X) and 500 μL of 12.5% wash buffer (12.5% formamide, 2×SSC and 0.1% Triton X-100). Another 500 μL of DAPI solution (4×SSCT solution with DAPI at a 1:100 dilution, Dojindo, 340-07971) was flowed and incubated for 30 seconds at room temperature. An anti-bleaching buffer (50 mM Tris-HCl pH 8.0, 300 mM NaCl, 2×SSC, 3 mM trolox (Sigma-Aldrich, 238813), 0.8% D-(+)-Glucose (Nacalai Tesque, 16806-25), 1,000-fold diluted catalase (Sigma-Aldrich, C3155), 0.5 mg/ml glucose oxidase (Sigma-Aldrich, G2133)) was then flowed through the system, followed by imaging 150 seconds later. Post-imaging, 1.4 mL of 55% wash buffer (55% formamide, 2×SSC and 0.1% Triton X-100) was applied for a 2-minute stripping operation at room temperature. The samples were subsequently washed twice with 4×SSCT. These procedures were repeated for each set of readout and secondary probes. In the final round of seq-RNA-FISH, images were captured using readouts against polyT LNA oligonucleotide to stain all RNA. After seq-RNA-FISH, samples were removed from the microscope for further experimental procedures preceding seq-DNA-FISH imaging (refer to the “seq-DNA/RNA/IF-FISH” section). For seq-DNA-FISH and seq-IF-FISH imaging, FOVs previously captured during seq-RNA-FISH were used. During seq-DNA-FISH, a mixture of secondary and readout probes for fiducial marker staining (comprising three types: Alexa488, ATTO565, and Alexa647) was each added at 50 nM. After seq-DNA-FISH imaging, to avoid contamination with fiducial markers, the samples underwent another stripping procedure, and the fluidics delivery system was cleaned. If significant shifts in imaging position occurred, additional rounds were executed. At the end of the day, corrections were made for any nuclear deformation and misalignments caused by fluidic delivery.

### Establishment of a cell line with deletion of the −190 kb region

In each well of 24-well gelatin-coated plates (Nunc), we plated 5 × 10^5^ C57BL6J ES cells with knocked-in *mNeonGreen* (*GFP*) and *iRFP670* (*iRFP*) in their respective *Nanog* alleles (referred to as NGiR) in 0.5 ml of 2i medium. Cells were cultured for 1 hour. Transfection complexes were prepared as follows: In one 1.5 ml tube, 25 µL of opti-MEM, 1 µL of P3000 reagent, 500 ng of eSpCas9-EF-49_10_L1, 500 ng of eSpCas9-EF-49_10_R1, and 300 ng of pCAG-mTagBFP2 (Addgene plasmid #122373) were combined. In another tube, 25 µL of opti-MEM and 1.8 µL of Lipofectamine 3000 were mixed well. The contents of both tubes were combined and incubated for 15 minutes before being added to the plated cells. The cells were cultured overnight, defining this day as Day 0. The medium was replaced on Day 1. On Day 2, an approximate 1000 BFP-positive cells, presumed to be successfully transfected, were sorted using FACSAria III cell sorter (BD Biosciences, Franklin Lakes, NJ, USA) and plated on a 6 cm dish. The medium was refreshed with new 2i medium on Day 4. On Day 7, 24 colonies were picked for downstream analysis and verification of gene targeting. PCR was performed using Δ-190_gPCR-F1 and Δ-190_gPCR-R1 primers. No cell lines with deletions on both alleles were obtained, thus candidate lines presumed to have deletion in a single allele were selected for further analysis. For additional verification, regions surrounding the upstream and downstream CRISPR target sequences were amplified and sequenced using the following primers: Δ-190_gPCR-F2 and Δ-190_gPCR-R3 for the upstream region, and Δ-190_gPCR-F3 and Δ-190_gPCR-R2 for the downstream region. Additionally, the deleted region was amplified and sequenced using Δ-190_gPCR-F2 and Δ-190_gPCR-R2 primers through Sanger sequencing. The linkage between the deleted −190kb region and the *Nanog-iRFP* knock-in allele was confirmed through single-molecule RNA fluorescence in situ hybridization (smRNA-FISH) using GFP/iRFP probes, as well as DNA-FISH employing a probe specific to the −190kb region. The confirmed cell line was designated as NG+iRΔ-190.

### smRNA-FISH

Both NGiR and NG+iRΔ-190 cell lines were seeded onto 8-well chambered cover glasses coated with LN511 at a density of 7.5×10^4^ cells per well in 2i media. The following day, cells were fixed by a 10-minute incubation with 4% PFA and subsequently washed three times with PBS. The samples were then permeabilized in 70% ethanol at −20°C for 1 hour and air-dried for 10 minutes. For cellular membrane disruption, samples were treated with 0.5% Triton-X in PBS at room temperature for 15 minutes and washed again three times with PBS. Cells were blocked for 15 minutes at room temperature using a blocking solution containing PBS, 10 mg/ml UltraPure BSA (Invitrogen AM2616), 0.3% Triton-X, 0.1% dextran sulfate (Nacalai Tesque, Kyoto, Japan, 03879-72), and 0.5 mg/ml sheared Salmon Sperm DNA (Invitrogen AM9680). Post-blocking, the samples were rinsed with 2×SSC (Nacalai Tesque, Kyoto, Japan, 32146-91). Probes for mNeonGreen (GFP) and iRFP670 conjugated with CAL Fluor Red 590 and Quasar 670 ^10^, along with 10 nM polyT LNA oligonucleotide, were hybridized in 30% hybridization buffer. The hybridization was carried out at 37°C for 12 hours in a humidified chamber. Following hybridization, samples were washed with a 35% wash buffer (35% formamide, 2×SSC and 0.1% Triton X-100) at room temperature for 30 minutes and subsequently rinsed three times with 4×SSC. The cells were then stained for 30 seconds at room temperature with a DAPI staining solution (4×SSCT solution with DAPI at 1:100 dilution; Dojindo, 340-07971). Finally, an anti-bleaching buffer containing 50 mM Tris-HCl pH 8.0, 300 mM NaCl, 2×SSC, 3 mM trolox (Sigma-Aldrich, 238813), 0.8% D-(+)- Glucose (Nacalai Tesque, Kyoto, Japan, 16806-25), 1,000-fold diluted catalase (Sigma-Aldrich, C3155), and 0.5 mg/ml glucose oxidase (Sigma-Aldrich, G2133) was added prior to imaging via microscopy.

### Pre-processing of seq-DNA/RNA/IF-FISH imaging data

Imaging data from seq-DNA/RNA/IF-FISH were initially processed using Fiji software. Subsequently, each field of view was chronologically organized along the time axis. To correct uneven background illumination, a dark image subtraction was performed. To prevent pixel values from becoming zero, a constant value of 1 was added to the entire image set. Flat-field correction was applied to each fluorescence image by dividing the images by the normalized background illumination while retaining the intensity profiles of fluorescent spots. Subsequently, drift correction was executed based on the DAPI images using ImageJ’s “Correct 3D drift” function (parameters: edge_enhance only=0, lowest=1, highest=51, max_shift_x=50, max_shift_y=50, max_shift_z=30). Hyper-stack images were subjected to minimum intensity projection along the z and time dimensions, followed by cropping to exclude xy regions containing zero values. Similarly, the hyper-stack images were processed in the x and time dimensions to ensure the exclusion of yz regions containing zero values. Additional drift correction was performed based on DAPI images using Fiji’s “Descriptor-based series registration (2d/3d + t)” (parameters: series_of_images=stack, brightness_of=Low, approximate_size=[5 px], type_of_detections=[Minima & Maxima], subpixel_localization=[3-dimensional quadratic fit], transformation_model=[Affine (3d)], images_are_roughly_aligned, number_of_neighbors=3, redundancy=1, significance=3, allowed_error_for_ransac=5, global_optimization=[All against first image (no global optimization)], range=5, choose_registration_channel=1, image=[Fuse and display], interpolation=[Linear Interpolation]). Henceforth, voxel dimensions are x:y:z = 130:130:130 nm. At this point, only seq-DNA-FISH images detected with a 647 nm readout probe were extracted and subjected to average intensity projection along the time axis. Given that all images contain fiducial markers, this procedure emphasizes these fiducial regions, saved as Fiducial_enhance_stack.tif. Subsequently, maximum intensity projection was applied along the time axis to the 647 nm seq-DNA-FISH images, thereby generating an overlaid image of the detected foci, saved as seqDNAFISH_647_foci_enhance_stack.tif. Post-drift correction images were saved separately for each field of view, capture timing, and color channel.

### Nuclear segmentation of seq-DNA/RNA/IF-FISH data

3D segmentation of the cell nuclei was performed using Cellpose (version 0.6.5)^42^. Initially, nuclear images were opened and resampled to reduce the xy resolution by a factor of 1/5. These images were further subjected to Gaussian blurring with a standard deviation of 1.5 pixels. Utilizing these preprocessed images, 3D segmentation was carried out using Cellpose. The original xy resolution was restored, and a series of morphological operations were executed: erosion with a radius of 1, dilation with a radius of 2, opening with a radius of 8, and another dilation with a radius of 4. These processed images were saved as regions of interest (ROIs). These provisional ROIs were then visually verified and further refined manually using Napari (DOI: 10.5281/zenodo.3555620). Nuclei touching the xy boundaries of the image were excluded from the analysis.

### seq-DNA-FISH analysis

For the detection of fluorescent spots in seq-DNA-FISH, we employed Big-FISH (version 0.6.2)^43^. Fluorescent spots were initially detected employing the ’detection.detect_spots’ function, with parameters set as follows: ’return_threshold=True,’ and ’voxel_size’ and ’spot_radius’ both set to (130, 130, 130) and (240, 189, 189), respectively. Subsequent to this initial detection, subpixel fitting was executed using the ’detection.fit_subpixel’ function, maintaining the same ’voxel_size’ and ’spot_radius’ parameters. It is noteworthy that any detected spots not falling within the nuclear regions as delineated in “Nuclear segmentation of seq-DNA/RNA/IF-FISH data” were excluded from further analysis. Moreover, any spots with fluorescence intensities ranking outside the top 10 within individual cells were also omitted from the analysis. Subsequently, Fiducial_enhance_stack.tif was utilized for spot identification via Big-FISH. All detected spots within a 3-pixel radius were considered as fiducial markers. In different channel images, fiducials in close proximity (within 2 pixels) were associated. However, if two fiducial candidate spots in the same channel were located within 3 pixels of each other, both were excluded from consideration as fiducial markers due to potential calibration difficulties. The 647 nm channel was considered the fiducial, and the offset between fiducial marker candidates in the same capture series across three channels was calculated. Using the chromatic offset information from the closest neighboring fiducial marker, other spot positions were adjusted. These spots were subsequently verified using Napari, and any that were unequivocally identified as fiducial markers were excluded. Spot intensities were normalized to the median nuclear intensity, and spots below a value of 1.1 were omitted from analysis. Subsequently, only the top four brightest spots within each channel and each cell during each capture were selected for analysis, excluding the rest. Further, we applied aligner.find_all_chr from Jie^44^ to assess whether the fluorescent spots were located on the same chromosome, employing parameters : cell_pts_input = cell_pts_input, gene_dist = gene_dist.values.astype(’int’), bin_size = bin_size, nm_per_bp = 0.34, pixel_dist= 130.0, num_skip = 5, total_num_skip_frac = 0.8, norm_skip_penalty = False, stretch_factor = 1.01, init_skip_frac = 0.15, lim_init_skip = False, max_iter = 2. For chromosomes appearing to have swapped connections in proximity, correction was performed by determining two centroids using k-means (KMeans(n_clusters=2)), and chromosomes were reassigned to the nearest centroid. From the assigned fluorescent spots, median xyz coordinates were calculated for each chromosome. Distances from these median coordinates to each spot were computed. Outliers exceeding the threshold of average distance + 2 × standard deviation (34.9 pixels) were excluded from the analysis. Target genomic regions with detection efficiency below half of the overall median detection efficiency were also excluded (Supplementary Fig. 3a). Furthermore, in the median distance matrix of Replicate 1, data corresponding to three regions that displayed patterns distinctly divergent from those of Replicate 2 were excluded from the analysis (Supplementary Fig. 3a).

### seq-RNA-FISH analysis

For the spot detection in preprocessed seq-RNA-FISH images, we employed Big-FISH. The analysis was conducted using the detection.detect_spots function with the parameters: return_threshold=True, voxel_size=(130, 130, 130), and spot_radius=(240, 189, 189). While the automatically determined threshold was often deemed sufficient, manual adjustments were made when necessary. Based on the set threshold, fluorescent spots were identified. Further, transcriptional foci were discerned utilizing the detection.decompose_dense and detection.detect_clusters functions with a minimum requirement of three spots (nb_min_spots=3). Subsequently, the presence or absence of these transcriptional foci was determined based on the coordinates of the gene vicinity as ascertained through seq-DNA-FISH analysis. If the distance from the median coordinates of the specific chromosome’s seq-DNA-FISH spots was within 30 pixels, transcription was considered to have occurred.

### seq-IF-FISH analysis

Utilizing the nuclear ROI determined through “Nuclear segmentation of seq-DNA/RNA/IF-FISH data,” we calculated the mean fluorescence intensity within cell nuclei. Only proteins/PTMs with a relative intensity value exceeding 1.5 when compared to the same channel’s negative control—captured without the addition of readout probe and secondary probe—were considered for analysis (Supplementary Fig. 5b,c). Notably, HDAC3 and SMAD4, which exhibited localization patterns divergent from those typically obtained through general immunofluorescence, were excluded from the study (Supplementary Fig. 5a,c).

In 1 Mb intervals, we examined the correlation between ChIP-seq enrichment levels and fluorescence intensity data obtained from seq-IF-FISH. Initially, 55 intervals of 1Mb each were designated starting at chr6:94,672,507. However, segments numbered 9, 11, 16, 18, and 36 were excluded as the corresponding seq-DNA-FISH data were ultimately not analyzed (Supplementary Fig. 3a). Using genomeCoverageBed, ChIP-seq enrichment levels were calculated in terms of reads per million (RPM). The pertinent Bigwig files are listed in Supplementary Table 7. Within the seq-IF-FISH images, the mean fluorescence intensity was calculated in a spherical region with a radius of 3 pixels, centered on the 3D coordinates of specific genome regions as determined by seq-DNA-FISH analysis. When multiple seq-DNA-FISH targets were included within the predefined segments, this mean value was employed. From these parameters, we calculated the Z-score as (Mean fluorescence intensity of the sphere - Mean nuclear intensity) / Standard deviation of nuclear intensity.

For the detection of fluorescent spots, Big-FISH was utilized. Spots were identified using the detection.detect_spots function with parameters: threshold=40, voxel_size=(130, 130, 130), and spot_radius=(200, 200, 200). Subpixel fitting was performed on these detected spots using the detection.fit_subpixel function, also with parameters: voxel_size=(130, 130, 130), and spot_radius=(200, 200, 200). The distance to the 3D coordinates of each target, as determined by seq-DNA-FISH analysis, was calculated. The entity closest to a specific genome coordinate was classified as the nearest condensates.

### A/B compartment calculation

First, we used the *.hic* formatted file (4DNFI4OUMWZ8 in the 4DN data portal) from *in situ* Hi-C experiments on mouse embryonic stem cells^19^. Next, using Straw^45^, we extracted the KR-normalized observed/expected matrix for chromosome 6 at 25-kb resolution. We then calculated the first eigenvector of the log2 ratio of the observed-to-expected matrix. Finally, we flipped the eigenvector profile to ensure a positive correlation coefficient with the gene density profile^46^.

### Micro-C data analysis

We used Micro-C data (4DNFI6HG4GP3 for WT in the 4DN data portal; GSE178982 for mouse ES cells with acute depletion of RAD21 and CTCF in the Gene Expression Omnibus repository) on JM8.N4 mouse ES cells^26,27^ to calculate the normalized contact probability matrices for the *Nanog* region (chr6: 122,350–122,900 kb) using 25-kb bins. For ΔCTCF and ΔRAD21 Micro-C data, we initially generated *.hic* files via the 4DN Hi-C data processing pipeline. Subsequently, using Straw^45^, we extracted the KR-normalized matrices. Additionally, we computed the normalized contact probability matrices using PHi-C2 *preprocessing* command so that the diagonal elements become the unit value^30,31^. We then compared these normalized contact probability matrices by calculating the log2 ratio of mouse ES cells with acute depletion of RAD21 and CTCF to JM8.N4 mouse ES cells.

### Modeling of higher-order genomic structural dynamics using PHi-C

Based on seq-DNA/RNA-FISH data, we can generate the contact probability map with the 350-nm distance threshold according to a specific gene’s transcriptionally active/inactive state. We analyzed the genomic regions of *Nanog* and *Sox2* from our data and MERFISH data^33^ with 25-kb and 5-kb resolutions, respectively. Then, we input the generated contact probability data into the PHi-C2 pipeline^31^. First, the PHi-C pipeline reconstructs a contact probability map through optimization of the parameters of the PHi-C polymer model. Here, we used the default input parameters of the PHi-C2 *optimization* command. The output contact probability maps were in good agreement with the inputs (Figs. 5a, 6a).

Then, the PHi-C pipeline allows for calculating structure and dynamics using the optimized parameters of the PHi-C polymer model. However, the spatial and temporal scales of the polymer model are normalized in the units. To recover the model in actual spatial and temporal scales, we need two parameters: the contact distance *σ* [m] and the friction coefficient of a polymer segment *γ* [kg/s]. We set *σ* = 350 [nm]. To obtain the value of *γ* in active and inactive states, we used the mean-square displacement (MSD) data of *Nanog* and *Sox2*^11^. First, we multiplied the MSD values by 1.5 to convert from 2-dimension to 3-dimension. Then, we fitted the value of *γ* using the following theoretical MSD of the PHi-C model^30^:

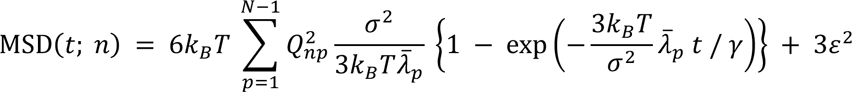

where *t* represents time, *k_B_* is the Boltzmann constant (= 1.380649×10^-23^ [m^2^ kg s^-2^ K^-1^]), *T* means the temperature set as 310 [K], *Q* is the orthogonal matrix of the optimized Laplacian matrix of the PHi-C polymer model, 𝜆̄_𝘱_ is the normalized eigenvalue of the *p*th mode, *n* is the matrix index of *Nanog* or *Sox2* region, and *ε* represents the estimation error in position determination^47^ and is also a fitted parameter. We used a SciPy function (scipy.optimize.curve_fit) in the fitting.

### Analysis of duration time between E-P regions in PHi-C dynamics simulations

We obtained polymer models consistent with structures and dynamics in actual spatial and temporal scales based on the above PHi-C modeling to the contact matrices from seq-DNA/RNA-FISH data. Then, we calculated the duration times for which E-P regions (*Nanog* and −45SE, 60SE, −190 kb; *Sox2* and SE) are close in PHi-C dynamics simulations (Figs. 5c-f, 6). In the PHi-C2 *dynamics* command, we set the stepsize parameter so that the integration timestep corresponds to actual time Δ*t* = 0.1 [s]: *k*_B_*T*Δ*t* / (*γσ*^2^), where *γ* is the fitted friction coefficient and *σ* = 350 [nm]. In addition, we numerically integrated 10^5^ steps corresponding to 10^4^ seconds, for 100 different initial conformations. Finally, we measured the duration times that E-P regions are in proximity threshold distances 100 [nm], 200 [nm], and 350 [nm]. First, we estimated the probability density of the duration times by the following two-component exponential model: 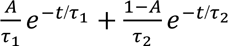. Here, *τ*_1_ and *τ*_2_ (> *τ*_1_) represent the characteristic duration times between E-P regions. Then, we calculated the average duration time by 𝐴𝜏_1_ + (1 − 𝐴)𝜏_2_.

### Visualization of PHi-C dynamics simulations

In the visualization (Supplementary Movies 2 and 3), we fixed the center of mass of polymer conformations to the origin, and calculated the dynamics and distance maps within 50 seconds. We used VMD to visualize polymer dynamics^48^.

### Statistics and Reproducibility and plots

The exact number, *n*, of data points and their representation (such as cells and independent experiments), and statistical tests used are indicated in the respective Figure legends and in the results. All experiments were performed as two or more independent experiments. The same conclusions were obtained from each experiment. Statistical tests were performed in R software (The R Project for Statistical Computing, Vienna, Austria) and Python (Python Software Foundation, https://www.python.org/).

## Data and code availability

All data are available in the main text or the supplementary materials. Publicly available software and packages were used in this study.

Any additional information required to reanalyze the data reported in this paper is available from the lead contact upon request.

## Acknowledgements

We thank Y. Ochiai (Kyushu Univ., Japan) and M. Shintani (Hiroshima Univ., Japan) for providing technical assistance. We extend our heartfelt gratitude to I. Hiratani (RIKEN, Japan) and H. Miura (RIKEN, Japan) for their invaluable discussions about seq-DNA-FISH analysis. We also thank L. Cai (Caltech) for providing insights into the automation of seq-DNA/RNA/IF-FISH, and S. Tsuda (Ansanga Lab) for their indispensable support in the fabrication and control of the automation apparatus. Additionally, we appreciate the critical feedback and insightful comments on the manuscript provided by M. Francois (University of Sydney) and I. Solovei (Ludwig-Maximilians-University of Munich), which greatly enriched the content of our work. H.Oc was supported by Grants-in Aid for Scientific Research from the Ministry of Education, Culture, Sports, Science, and Technology (grant nos. JP21H05753, JP22H02609, and JP22H04694), and NIG-JOINT (grant nos. 76A2023 and 15R2023). H.Oh was supported by Grants-in Aid for Scientific Research from the Ministry of Education, Culture, Sports, Science, and Technology (grant no. JP22K15084). Y.O was supported by Grants-in Aid for Scientific Research from the Ministry of Education, Culture, Sports, Science, and Technology (grant no. JP18H05527). S.S was supported by Grants-in Aid for Scientific Research from the Ministry of Education, Culture, Sports, Science, and Technology (grant no. JP23H04297). H.Oc, H. Oh and Y.O. are also supported by the Medical Research Center Initiative for High Depth Omics at Kyushu University, Japan. H.Oc. and S.S. are also supported by the RIKEN-Hiroshima University Science and Technology Hub Collaborative Research Program.

## Author contributions

H.Oc. conceived and supervised the project. H.Oc., H.Oh., and S.S. designed the experiments. H.Oh., S.S., H.Oc., H.Ow., T.F., K.H., S.O., T.Y., and Y.O. performed the experiments. H.Oh., S.S. and H.Oc. analyzed the data. H.Oc. H.Oh., S.S., Y.O., K.H., wrote the paper.

## Competing interests

Authors declare that they have no competing interests.

## Materials & Correspondence

Correspondence and requests for materials should be addressed to Hiroshi Ochiai.

## Supplementary information

**Supplementary Fig. 1.**
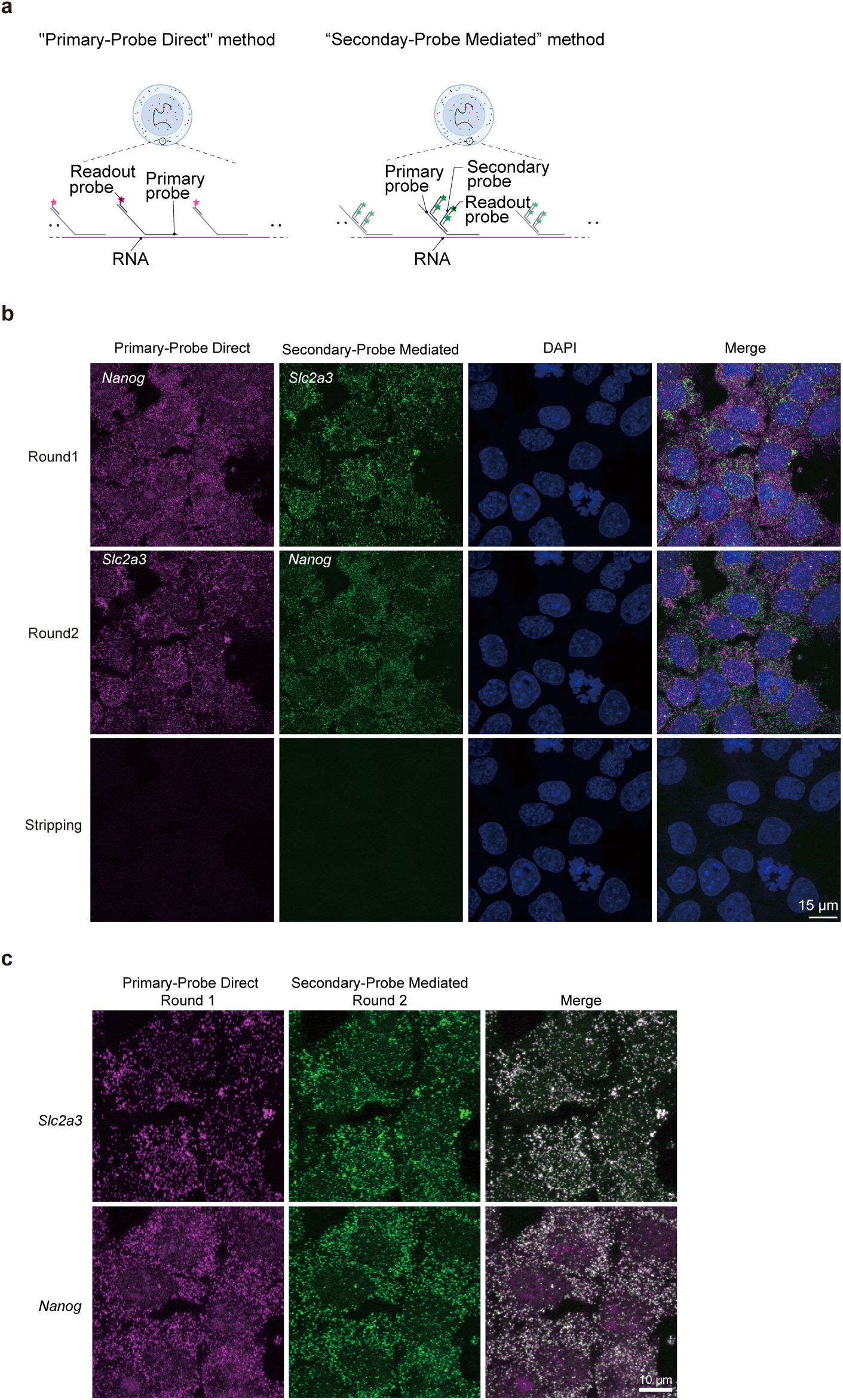
Validation of spot detection method in seq-RNA-FISH. **a** Schematic representation of the fluorescent labeling strategies employed in seq-DNA/RNA/IF-FISH, specifically the “Primary-Probe Direct” and “Secondary-Probe Mediated” methods, with an example focusing on RNA detection. In the “Primary-Probe Direct” detection method, a primary probe equipped with an adapter sequence capable of binding a readout probe is first hybridized to the target molecule, in this case, RNA. Subsequently, the fluorescence-tagged readout probe is hybridized, rendering the target molecule visible. Synthesizing these specialized readout probes often requires additional purification methods such as High-Performance Liquid Chromatography (HPLC). Given the excessive cost associated with this, adopting this approach presents a significant hurdle. Conversely, the “Secondary-Probe Mediated” method introduces a secondary probe between the primary and readout probes. This design obviates the need for multiple unique readout probes, thereby facilitating their recycling for multiple targets. Although this approach necessitates the preparation of numerous non-fluorescent secondary probes, the readout probes need only be prepared in quantities corresponding to the number of fluorescent channels used (three types in our case: Alexa488, ATTO565, and Alexa647). We chose to employ secondary probes that were designed with sequences complementary to both the adapter sequences on the primary probes and specific sequences on the readout probes. By simultaneously adding and hybridizing both the secondary and readout probes to the cellular samples, the readout probes become localized at the site of the primary probes. This strategy requires preparing readout probes corresponding to the number of fluorescent dye types utilized, thereby streamlining the overall process and reducing cost. **b** Validation of both “Primary-Probe Direct” and “Secondary-Probe Mediated” seq-RNA-FISH methodologies. C57BL6/J mESCs cultured in 2i medium were subjected to seq-RNA-FISH. We designed either 24 or 48 primary probes corresponding to the “Primary-Probe Direct” or “Secondary-Probe Mediated” seq-RNA-FISH methods for the detection of *Nanog* and *Slc2a3* mRNA. These primary probes were initially hybridized. Subsequently, the appropriate readout and secondary probes were hybridized to visualize the RNA (Round 1). Following imaging via microscopy, the secondary and readout probes were stripped off, and a different set of suitable readout and secondary probes were hybridized to re-visualize the RNA (Round 2). Signals were observed using both “Primary-Probe Direct” labeling via readout probes and “Secondary-Probe Mediated” labeling via secondary probes. Finally, imaging was conducted after stripping the readout and secondary probes to confirm their successful removal as intended. **c** Fluorescent spots labeled by the “Primary-Probe Direct” method were found to match fluorescent spots labeled by the “Secondary-Probe Mediated” method.

**Supplementary Fig. 2.**
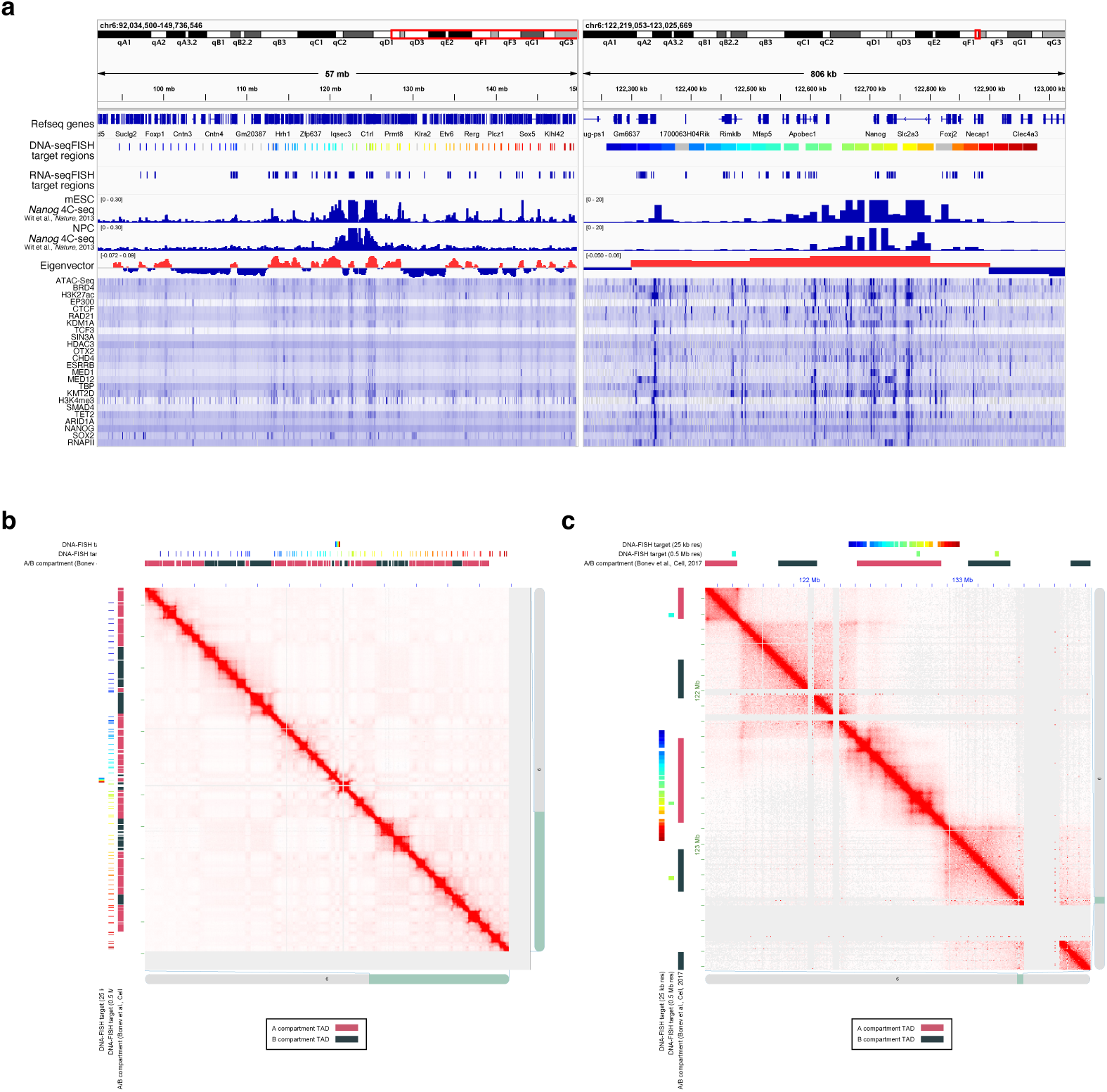
Characteristics of target regions for seq-DNA/RNA-FISH analysis in this study. **a** Target RNA, genomic regions, and corresponding epigenomic information in seq-DNA/RNA-FISH analysis. **b-c** High-resolution Hi-C data of the chromosome 6 region targeted in this study for seq-DNA-FISH^19^. Hi-C data within genomic regions where probes were designed at approximately 0.5 Mb intervals (**b**) or 25 kb intervals (**c**). The A/B compartments are denoted as annotated in Bonev et al.^19^. The regions designed at 25 kb intervals roughly correspond to a single Topologically Associating Domain (TAD).

**Supplementary Fig. 3.**
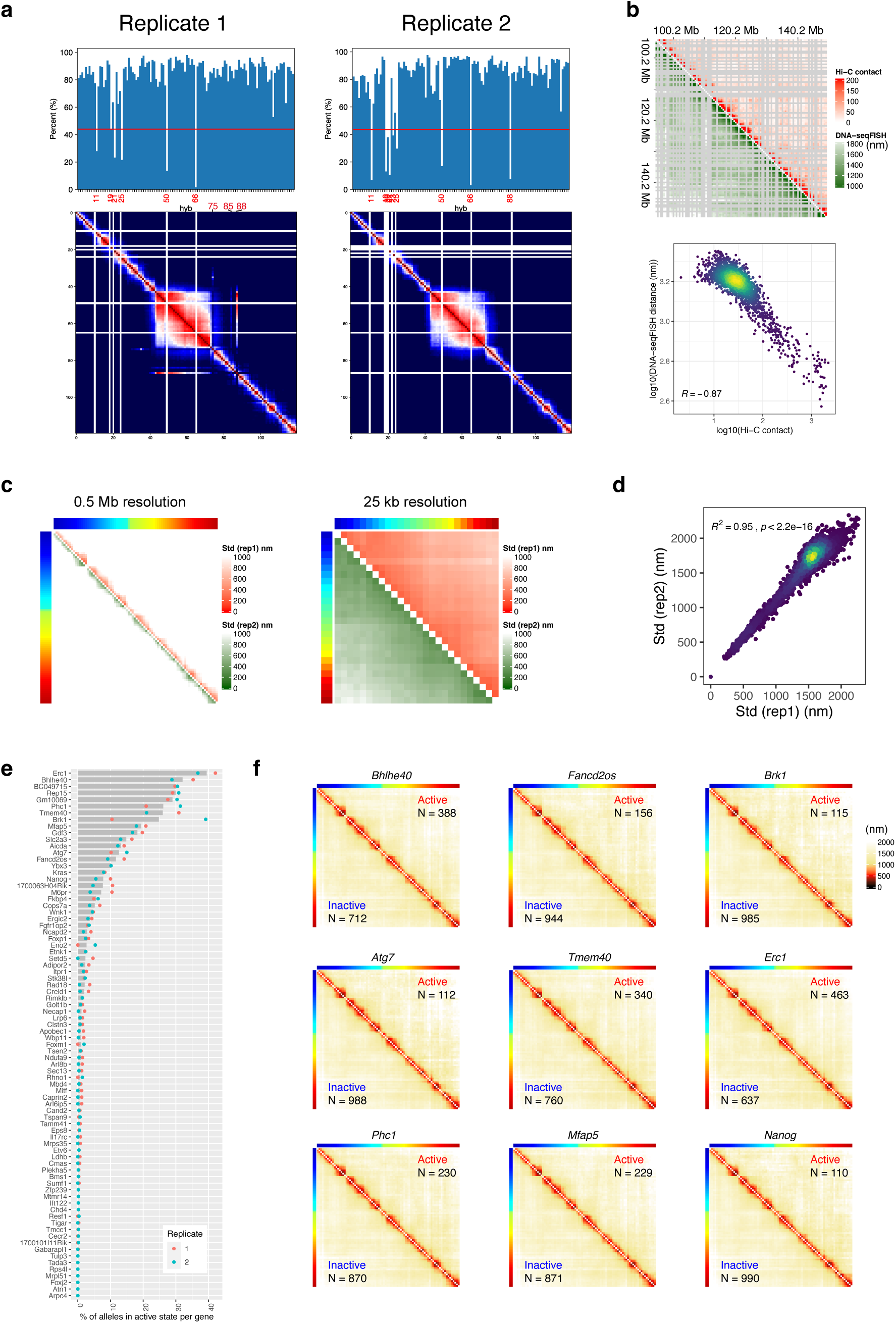
Quality check of seq-DNA-FISH experiments. **a** Efficacy of fluorescence signal detection for each targeted genomic region (Upper Panel) and corresponding median distance matrices in seq-DNA-FISH experiments (Lower Panel). The red line on the upper panel represents half of the average detection efficiency across all targeted regions in each experiment. Regions falling below this threshold were excluded from further analysis. In the median distance matrices, regions that are proximal in terms of 1D genomic distance are anticipated to exhibit spatial proximity in the 3D median distance as well. However, regions ID 74, 84, 87 in Replicate 1 deviated from this intuition and substantially differed from Replicate 2 data; thus, they were also excluded from the analysis. Replicate 1: *N* = 1,100 alleles; Replicate 2: *N* = 580 alleles. Unless otherwise stated, data from Replicate 1 are depicted in the subsequent Figures. **b** Comparison between Hi-C contact frequency^19^ and seq-DNA-FISH-derived median distance matrices, demonstrated at a 0.5 Mb resolution. Grey areas indicate regions not included in this analysis. The lower panel shows scatter plots of data from the upper panel. seq-DNA-FISH : *N* = 1,100. **c** Comparative median distance matrices acquired from seq-DNA-FISH in Replicates 1 and 2, depicted separately for 0.5 Mb and 25 kb resolutions. Replicate 1: *N* = 1,100; Replicate 2: *N* = 580. The color codes at the top and left edges represent targeted genomic regions from Fig. 1b. **d** Scatter plot comparison of data from **c**, incorporating both 25 kb and 0.5 Mb resolution data. High correlation coefficients are observed, indicating excellent reproducibility. **e** Proportion of cells displaying transcription foci. The bar graph represents the average percentage of cells with transcription foci in Replicates 1 and 2. **f** Median distance matrices for probe regions designed at 0.5 Mb intervals. The sample sizes are noted in the Figure. Data are partitioned by the active/inactive state of specific genes, and their respective median distance matrices are presented. The color codes at the top and left edges represent targeted genomic regions from Fig. 1b.

**Supplementary Fig. 4.**
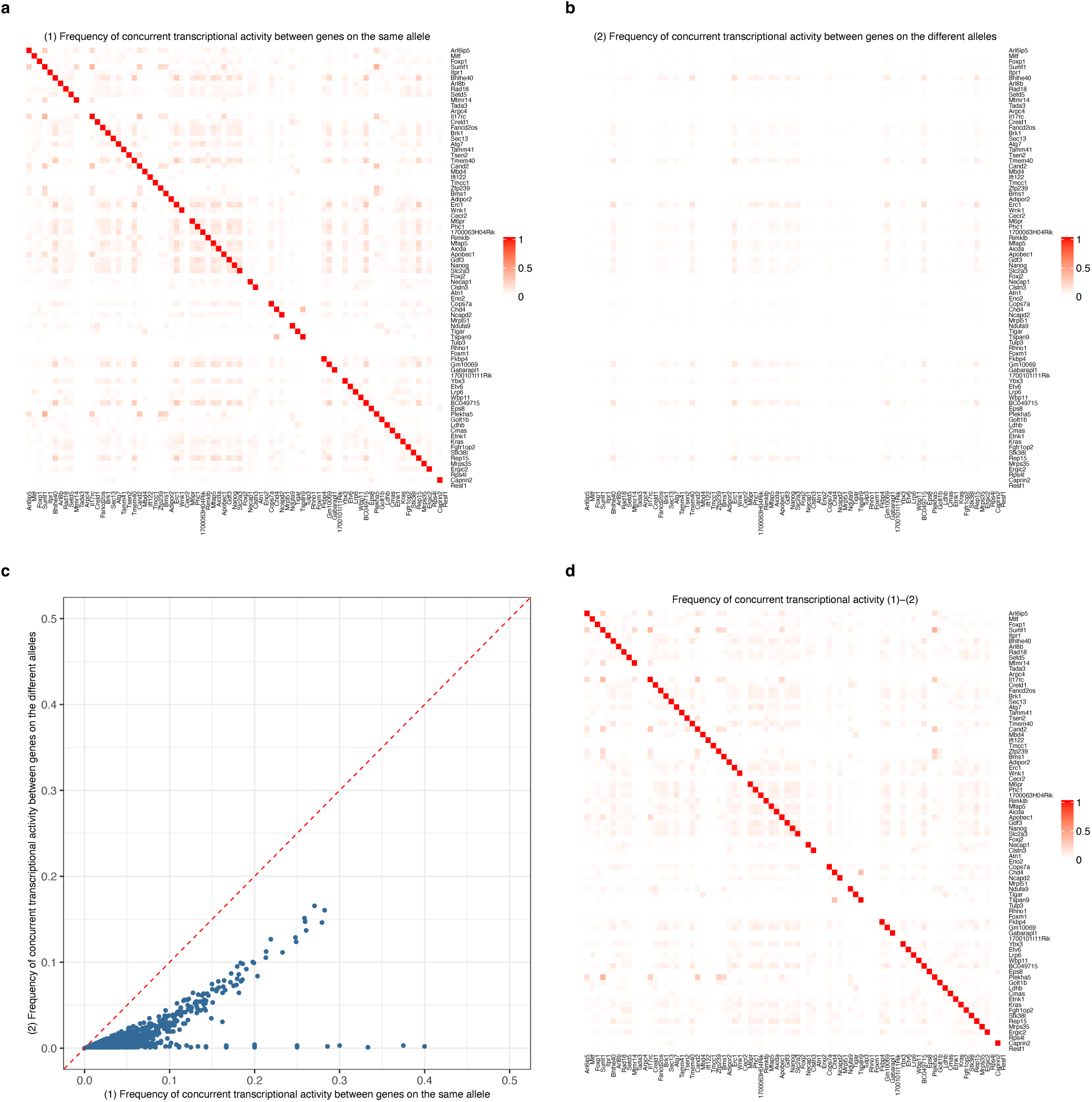
Frequency of concurrent transcriptional activity between genes on the same allele. **a** Data (1): Frequency of concurrent transcriptional activity between genes located on the same allele. The dataset comprises information from 1,100 alleles. Concurrent transcriptional activity is defined as the number of alleles where *Gene X* and *Gene Y* are simultaneously in the active state divided by the number of alleles where either *Gene X* or *Gene Y* is in the active state. Due to varying transcriptional frequencies for each gene, the sample size (*N*) differs between elements. **b** Data (2): Frequency of concurrent transcriptional activity between genes located on different alleles within the same cell. The dataset includes data from 489 cells. Concurrent transcriptional activity here is defined as the number of cells where *Gene X* on Allele 1 and *Gene Y* on Allele 2 are simultaneously in the active state, divided by the number of cells where either *Gene X* on Allele 1 or *Gene Y* on Allele 2 is in the active state. As with **a**, due to different transcriptional frequencies for each gene, the sample size (*N*) varies between elements. **c** Scatter plot of Data (1) and Data (2). The red line represents the *y* = *x* line. Genes that exhibit high frequency of concurrent transcriptional activity within the same allele also tend to have a high frequency of concurrent transcriptional activity between different alleles within the same cell. However, the frequency of concurrent transcriptional activity is markedly higher when both genes locate on the same allele. This suggests that while global nuclear factors (e.g., expression levels of transcription factors) might have a moderate influence on concurrent transcriptional activity, the genomic structure of locating on the same allele could partially induce concurrent transcriptional activity. **d** Heatmap of Data (1) -Data (2), derived from **a** and **b**. This reveals what could be termed the apparent true frequency of concurrent transcriptional activity on the same allele, accounting for the influence of overall nuclear factors that contribute to concurrent transcriptional activity. It is important to note that this calculation is a simplified model and may not fully capture the complexities involved. Additional, more detailed analyses are necessary to comprehensively understand the true nature of concurrent transcriptional activity on the same allele. Several factors, such as varying transcriptional frequencies, sample size differences, and the influence of global nuclear factors, may introduce biases or limitations in this estimate.

**Supplementary Fig. 5.**
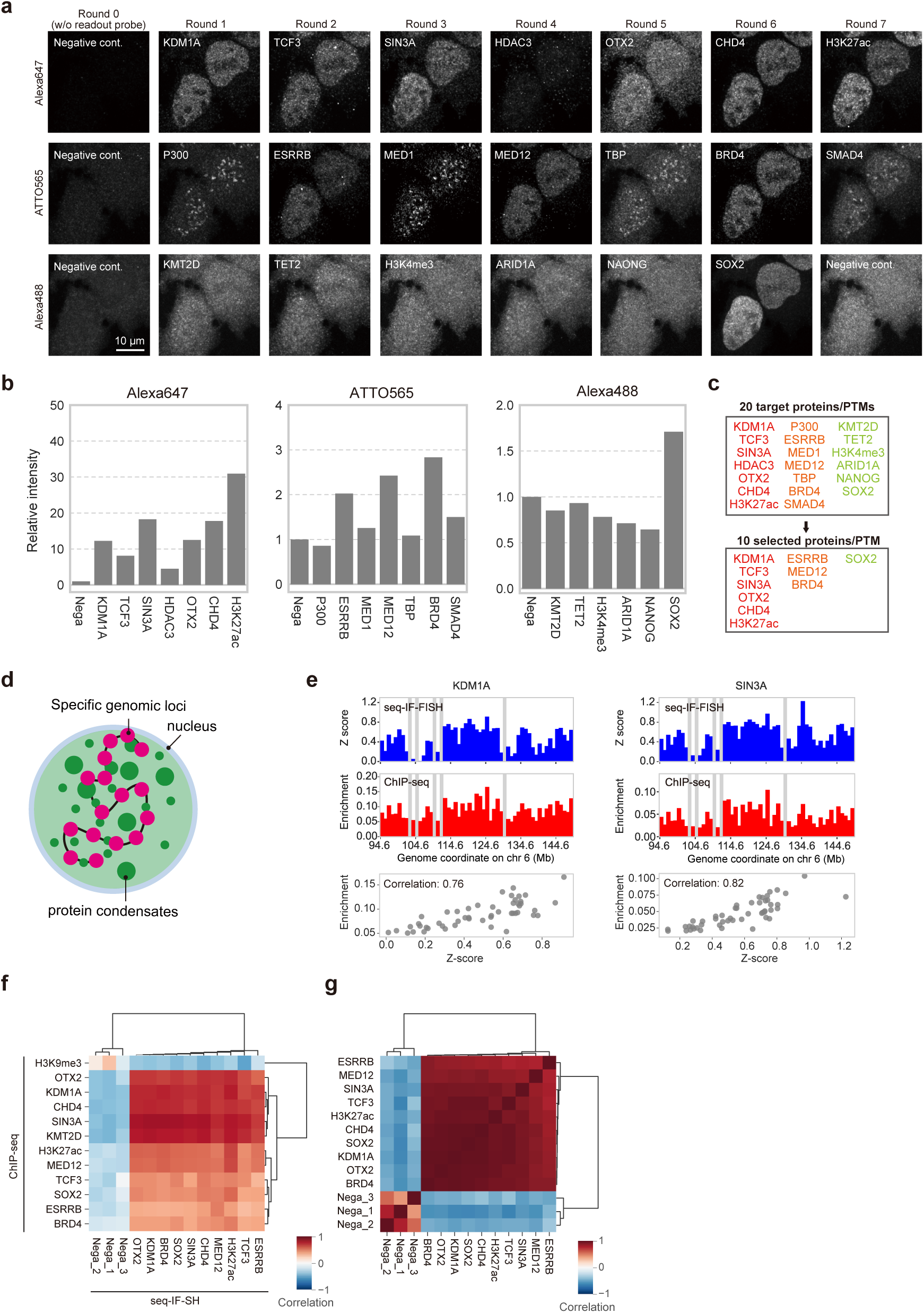
Quality check of seq-IF-FISH data. **a** Representative visualization of intranuclear proteins and post-translational modifications in seq-IF-FISH analysis. **b** Fluorescence intensity values were measured within the cell nucleus for each probe utilized in seq-IF-FISH. These values are presented as relative fluorescent intensities when compared to a negative control within the same fluorescence channel. Sample size: *N* = 611. **c** Initially targeted proteins and post-translational modifications (PTMs) in seq-IF-FISH are listed, along with those ultimately selected for further analysis. Proteins/PTMs with a relative fluorescent intensity value of 1.5 or below in **b** were excluded from further analysis. Additionally, HDAC3 and SMAD4, whose localization patterns substantially deviated from those obtained through general immunofluorescence, were also excluded. **d** Schematic representation of the localization of proteins/PTMs in specific genomic regions. **e** Correlation between fluorescence intensity data from seq-IF-FISH and ChIP-seq enrichment data, featuring KDM1A and SIN3A. The top graph displays fluorescent intensity Z-scores of KDM1A and SIN3A in seq-IF-FISH at specific genomic coordinates as determined by seq-DNA-FISH (details refer to Methods). *N* = 611. The middle graph represents ChIP-seq enrichment values for the same region (details refer to Methods). Gray areas denote locations excluded from analysis in seq-DNA-FISH. The bottom graph shows the scatter plot of fluorescent intensity Z-scores in seq-IF-FISH and ChIP-seq enrichment. **f** Heatmap of correlation values between fluorescent intensity Z-scores from seq-IF-FISH and ChIP-seq enrichment. “Nega_1,” “Nega_2,” and “Nega_3” represent the negative controls for the Alexa647, ATTO565, and Alexa488 fluorescent channels, respectively. **g** Heatmap of correlation values between fluorescent intensity Z-scores in seq-IF-FISH.

**Supplementary Fig. 6.**
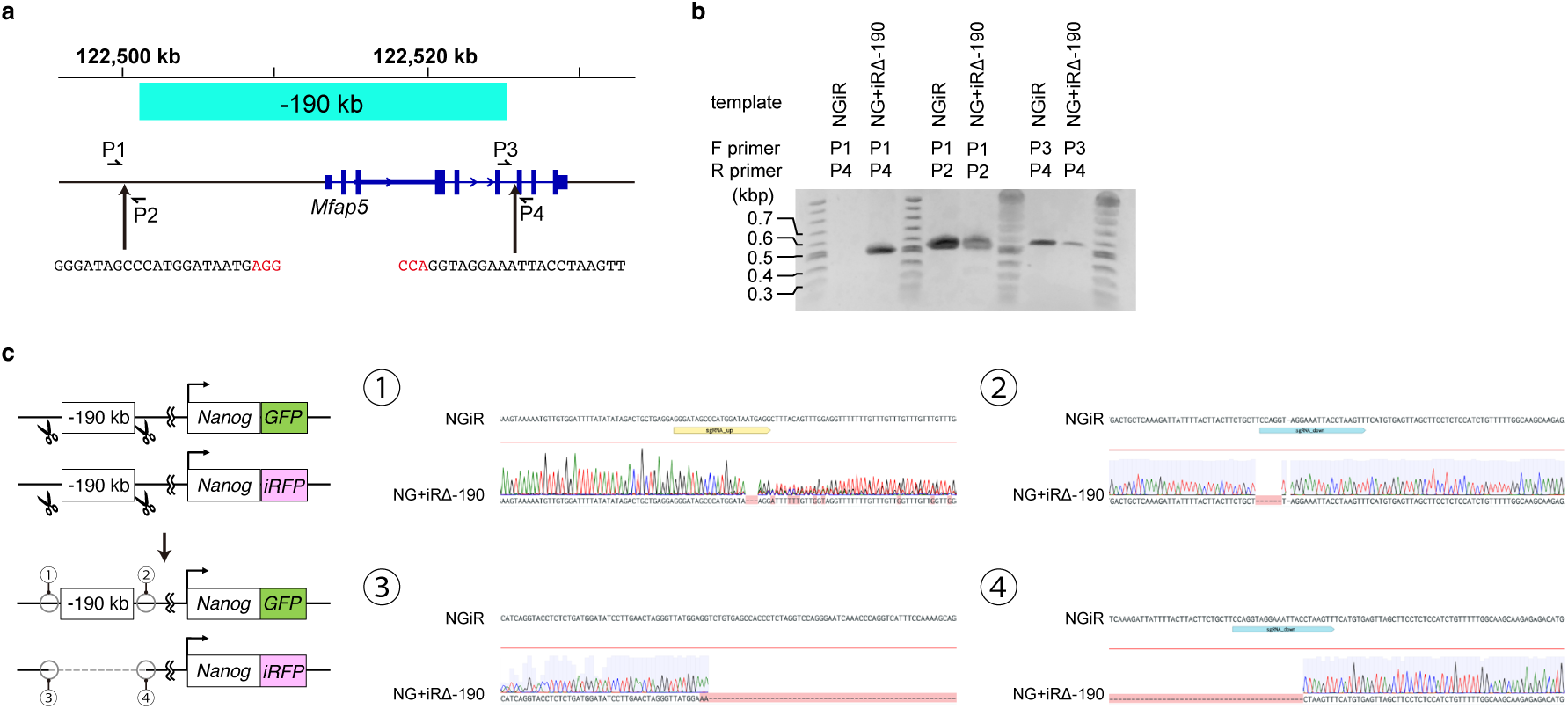
Establishment of a cell line with deletion of the −190 kb region. **a** Schematic representation of the −190 kb region, which encompasses the transcriptional start site of *Mfap5*. The nucleotide sequences noted at the bottom of the Figure represent the CRISPR target sites, with the Protospacer Adjacent Motif (PAM) sequence indicated in red. Positions P1 to P4 denote the locations of the primers used in **b**. **b** Genomic PCR of WT(NGiR) and deletion cell lines (NG+iRΔ-190). Genomic PCR was performed on both NGiR and NG+iRΔ-190, using PCR conditions optimized to allow amplification of sequences up to approximately 2 kb in length. Various combinations of the primers specified in **a** were used to perform the genomic PCR. In the out-out (P1 and P4 combination) primer set, amplification occurs in the NG+iRΔ-190 but not in the NGiR. However, amplification is observed in the NG+iRΔ-190 for both the P1-P2 and P3-P4 primer combinations, suggesting that only one allele is deleted in the NG+iRΔ-190 cell line. **c** Nucleotide sequence determination of the PCR-amplified fragments from genomic PCR. A deletion of 6 bases is observed in the downstream CRISPR target sequence of the non-deleted allele of NG+iRΔ-190. The upstream CRISPR target sequence of the non-deleted allele in NG+iRΔ-190 is unclear due to an ambiguous sequence chromatogram, although a deletion of a few bases is suspected. For the deleted allele of NG+iRΔ-190, a loss extends from the CRISPR target sequence up to approximately 190 bp upstream.

**Supplementary Fig. 7.**
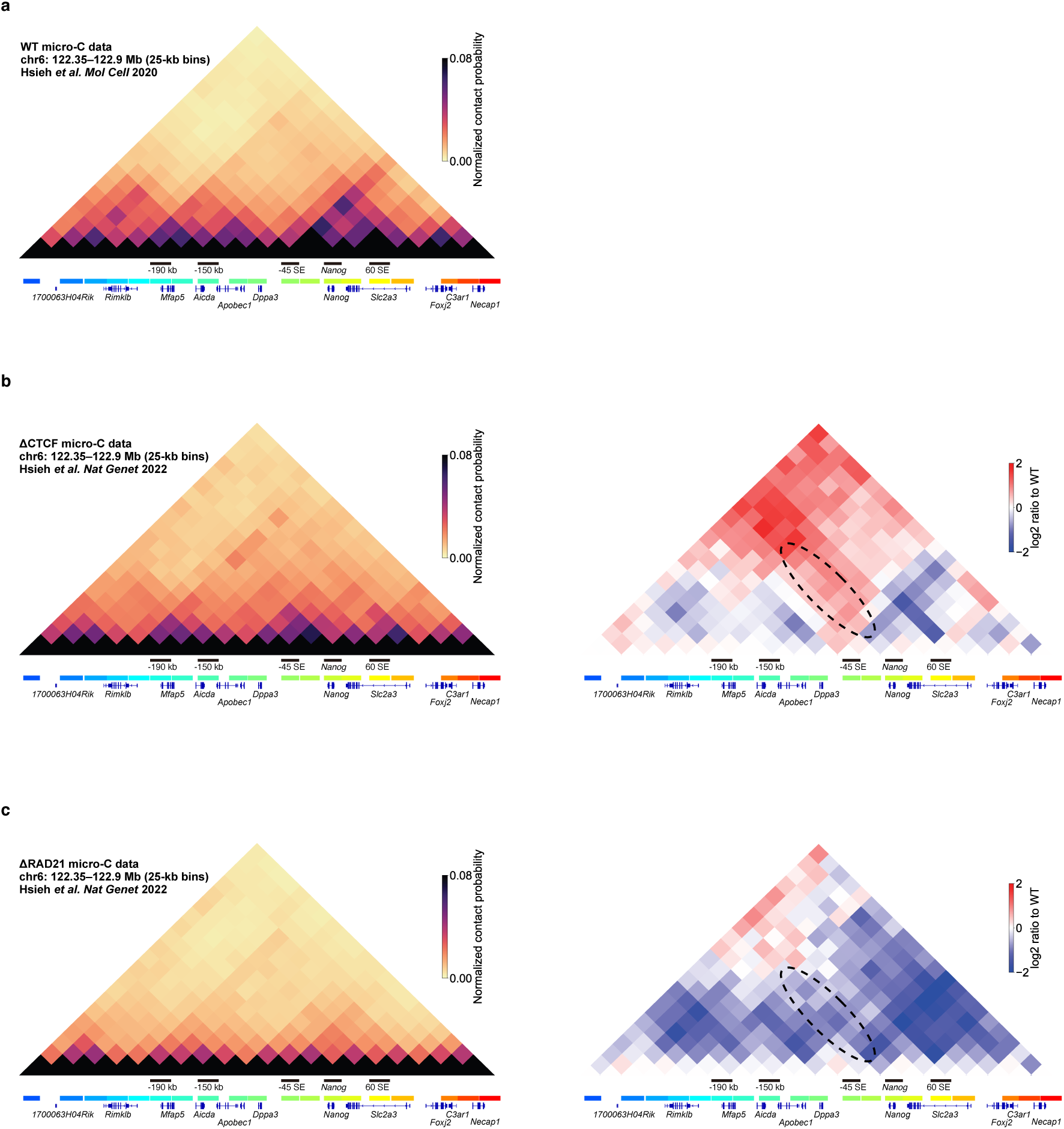
Micro-C data in mouse ES cells (25 kb resolution). **a** Micro-C data from WT mouse embryonic stem cells^26^. Contact frequencies have been normalized using PHi-C (see Methods for detail). **b** Micro-C data from mouse ES cells following acute degradation of CTCF^27^ (left panel). Contact frequencies are normalized using PHi-C (see Methods for detail). The right panel shows the relative contact frequencies in acute CTCF-degraded data compared to WT Micro-C data. Dotted ellipses indicate regions where contact frequency has increased in interaction domains with *Nanog* compared to the WT. **c** Micro-C data from mouse ES cells following acute degradation of RAD21, a subunit of cohesin^27^ (left panel). Contact frequencies are normalized using PHi-C (see Methods for detail). The right panel displays the relative contact frequencies in acute RAD21-degraded data compared to WT Micro-C data. Dotted ellipses indicate regions where contact frequency has decreased in interaction domains with *Nanog* compared to the WT.

## Supplementary information

**Supplementary Table 1: Oligonucleotide sequences used in this study.**

**Supplementary Table 2. Oligonucleotides used in the evaluation of the fluorescent spot detection method in seq-RNA-FISH.**

**Supplementary Table 3. Genomic regions targeted by seq-DNA-FISH analysis in this study.**

**Supplementary Table 4. Complementary sequences of target-binding sites for primary probes used in the seq-DNA-FISH analysis in this study.**

**Supplementary Table 5. Complementary sequences of target-binding sites for primary probes used in the seq-RNA-FISH analysis in this study.**

**Supplementary Table 6. List of antibodies used in seq-IF-FISH in this study.**

**Supplementary Table 7. List of public ChIP-seq data used in seq-IF-FISH quality check.**

**Supplementary Movie 1. Representative images of seq-DNA/RNA/IF-FISH.**

The movie is generated from maximum intensity projections of images obtained in each round using seq-DNA/RNA-IF-FISH. In each round, three types of both secondary and readout probe sets were used, enabling the acquisition of images across three channels. Each channel was utilized for observing the localization of a singular RNA, genomic region, protein, or post-translational modification. Another channel was designated for capturing images of DAPI-stained nuclei.

**Supplementary Movie 2. Genome dynamics in *Nanog* active state simulated by PHi-C.**

The movie illustrates a simulation of the dynamic genomic structure of the *Nanog* active state, based on seq-DNA/RNA-FISH data and modeled using PHi-C. The left panel displays the genomic structural dynamics, while the right panel presents a real-time inter-region distance matrix corresponding to that structure. In the active state, the genome exhibits elevated viscosity, resulting in relatively slow motion.

**Supplementary Movie 3. Genome dynamics in *Nanog* inactive state simulated by PHi-C.**

The movie illustrates a simulation of the dynamic genomic structure of the *Nanog* inactive state, leveraging seq-DNA/RNA-FISH data and modeled through PHi-C. The left panel reveals the dynamic genomic architecture, while the right panel displays a real-time inter-region distance matrix corresponding to that architecture.

Characteristically, in the inactive state, the genomic viscosity is lower compared to the active state, resulting in relatively rapid motion.

## Notes

### Competing Interest Statement

The authors have declared no competing interest.

